# Lipid droplets modulate proteostasis, SQST-1/SQSTM1 dynamics, and lifespan in *C. elegans*

**DOI:** 10.1101/2021.04.22.440991

**Authors:** Anita V. Kumar, Joslyn Mills, Wesley M. Parker, Joshua A. Leitão, Diego I. Rodriguez, Celeste Ng, Rishi Patel, Joseph L. Aguilera, Joseph R. Johnson, Shi Quan Wong, Louis R. Lapierre

## Abstract

The ability of organisms to live long depends largely on the maintenance of proteome stability via proteostatic mechanisms including translational regulation, protein chaperoning and degradation machineries. In several long-lived *Caenorhabditis elegans* strains, such as insulin/IGF-1 receptor *daf-2* mutants, enhanced proteostatic mechanisms are accompanied by elevated intestinal lipid stores, but the role of lipid droplets in longevity has remained obscure. Here, while determining the regulatory network of the selective autophagy receptor SQST-1/SQSTM1, we unexpectedly uncovered a novel role for lipid droplets in proteostasis and longevity. Using an unbiased genome-wide RNAi screening approach, we identified several SQST-1 modulators, including proteins found on lipid droplets and those prone to aggregate with age. SQST-1 accumulated on lipid droplets when autophagy was inhibited, suggesting that lipid droplets may serve a role in facilitating selective autophagy. Expansion of intestinal lipid droplets by silencing the conserved cytosolic triacylglycerol lipase gene *atgl-1/ATGL* enhanced autophagy, and extended lifespan in an HSF-1/HSF1-dependent and CDC-48/VCP-dependent manner. Silencing *atgl-1* mitigated the age-related accumulation of SQST-1 and reduced overall ubiquitination of proteins. Reducing *atgl-1* also improved proteostasis in proteotoxicity models of neurodegenerative diseases. Subcellular analyses revealed that lipid droplets unexpectedly harbor more soluble ubiquitinated proteins than the cytosol. Accordingly, low lipid droplet levels exacerbated the proteostatic collapse when autophagy or proteasome function was compromised. Altogether, our study uncovers a key role for lipid droplets in *C. elegans* as a proteostatic mediator that reduces protein ubiquitination, facilitates autophagy, and promotes longevity.

## INTRODUCTION

One of the major hallmarks of aging is the accumulation of damaged proteins that progressively compartmentalize in inclusions and aggregates^1^. In the nematode *C. elegans*, numerous proteins display impaired solubility with age^2,3^. This phenomenon suggests that mechanisms that mediate protein stabilization, such as chaperone-mediated folding and post translational modifications, actively contribute to somatic maintenance by preventing the collapse of the proteome^4,5^. The enhanced proteasomal^6^ and autophagic^7,8^ capacities, the increased chaperone function^9^ and alterations in ribosomal biogenesis^10^ and function^11,12^ in long-lived nematodes support the notion that the balance between synthesis, folding, and efficient clearance of proteins is important for conferring organismal longevity.

Generally, proteins destined for degradation are tagged via poly-ubiquitination and recognized by ubiquitin binding domain-containing proteins that direct cargo toward proteasomal or autophagic degradation. The autophagy receptor, Sequestosome 1 (SQST-1/SQSTM1) is a well-established and conserved mediator of cargo recognition, which includes ubiquitinated targets^13^. The SQST-1-cargo complex can interact with the autophagosome proteins LGG-1/GABARAP and LGG-2/LC3 to enable cargo sequestration in the nascent autophagosome^14^. Notably, SQST-1/SQSTM1 has emerged as a potential lifespan modulator in nematodes and flies^15-17^. Additionally, selective autophagy of mitochondria has been linked to longevity^18^, suggesting that autophagic degradation of specific cargos and organelles is beneficial against organismal aging. Although SQST-1/SQSTM1 has been implicated in the selective clearance of aggregated proteins^19^, much remains to be understood about the modulation of this autophagy receptor, its cargoes, and the nature and extent of its contribution to lifespan.

In nematodes, intestinal cells have a unique role in autophagy-mediated longevity as they manage nutrient influx and signals from neurons^20^ and the germline^21^ to coordinate appropriate responses necessary for organismal survival. In particular, intestinal autophagy genes are required for lifespan extension^22^ by maintaining intestinal tissue integrity and proteostasis. Intestinal cells are also sites of dynamic lysosomal function^23-26^ that mediate specific longevity-associated lipid signaling^27-29^. Intriguingly, several established long-lived animals including *daf-2, glp-1* and *rsks-1* mutants also maintain unusually large intestinal lipid droplet stores throughout their life^30-33^. Accordingly, redistributing lipids intended for secreted lipoproteins to intestinal lipid droplets is sufficient to extend lifespan^29^. However, in long-lived animals, it is unclear whether adult intestinal lipid stores are simply a consequence of reduced demand from the germline^34^, a long-term energy store rationing strategy^35^ or a more direct mediator of somatic maintenance. Here, while identifying the regulatory network of SQST-1, we found that lipid droplet accumulation in long-lived nematodes represents a novel proteostatic mechanism that reduces overall SQST-1 and ubiquitination levels, and mitigates age-related proteostatic stress.

## RESULTS

### SQST-1 over-expression is detrimental to lifespan in C. elegans

We initially considered the possibility that increasing SQST-1 may improve proteostasis by enhancing the ability of cells to mediate selective autophagy. We hypothesized that increasing the expression of a receptor that recognizes ubiquitinated cargoes may suffice in driving their degradation. While this study was being conducted, the Hansen laboratory showed that over-expressing SQST-1 can extend lifespan at 20°C^16^. Since longevity interventions and temperature mechanistically interact^36^, we examined lifespan at both 20°C and 25°C using several over-expressing strains (*psqst-1::sqst-1::rfp, psqst-1::sqst-1::gfp::rfp*), including two created in the Hansen laboratory (*psqst-1::sqst-1::gfp* and *psqst-1::sqst-1*)^16^. Over-expression of SQST-1 was unexpectedly detrimental at 25°C (**Figure 1a-c, Supplemental Figure 1a-c**) and was not sufficient to extend lifespan at 20°C (**Figure 1d-f**). While our findings at 20°C differ from recent literature^16^, highlighting potential lab-to-lab experimental variations, our findings are in line with most studies about the autophagy pathway and aging, i.e. that over-expressing an autophagy receptor might not necessarily be sufficient to improve lifespan^37^. At 25°C, SQST-1 clearly accumulates in the head, as well as the gonadal and intestinal tissues (**Supplemental Figure 1d**) as previously reported^38^. A closer investigation into the temperature-dependent differences in lifespan revealed marked upregulation of *sqst-1* mRNA at higher temperature in these strains (from ∼5-fold in wild-type, up to ∼75 fold in SQST-1::GFP over-expressing animals) (**Figure 1g**). In order to measure the active trafficking of SQST-1 into the lysosome as a guide for selective autophagy, we developed a tandem reporter system expressing SQST-1 fused to both GFP and RFP^7^ (GFP signal is quenched in low pH environments). We confirmed that the conversion of tandem SQST-1 into an RFP-only signal is reduced in *lgg-1-* or *lgg-2*-silenced animals (**Supplemental Figure 1e**). and autophagy-deficient *atg-18* mutants (**Supplemental Figure 1f**). While relatively modest at 20°C (**Figure 1h-i**), increasing temperature up to 30°C for 24 hours significantly enhanced the conversion to the RFP-only SQST-1 signal in the SQST-1 tandem reporter, suggesting that selective autophagy is induced with heat stress. Substantial temperature-dependent intestinal and neuronal accumulation of SQST-1 occurred during aging and over-expressing SQST-1 at 25°C was also accompanied by higher levels of total ubiquitinated proteins (**Figure 1a-f inset and Supplemental Figure 1g-h**). While over-expressing SQST-1 was detrimental at 25°C, so was the loss of *sqst-1* (**Supplemental Figure 1i-j**), suggesting that SQST-1 function has a temperature-specific role in the lifespan of wild-type animals. Expression and steady-state levels of SQST-1 likely need to be tightly coordinated with cargo targeting and more importantly, with the rate of protein degradation systems (i.e. autophagy and proteasome). Altogether, we found that solely increasing SQST-1 is detrimental for lifespan at 25°C and inconsequential at 20°C, indicating a temperature-dependent effect on SQST-1 dynamics in aging.

**Figure 1.**
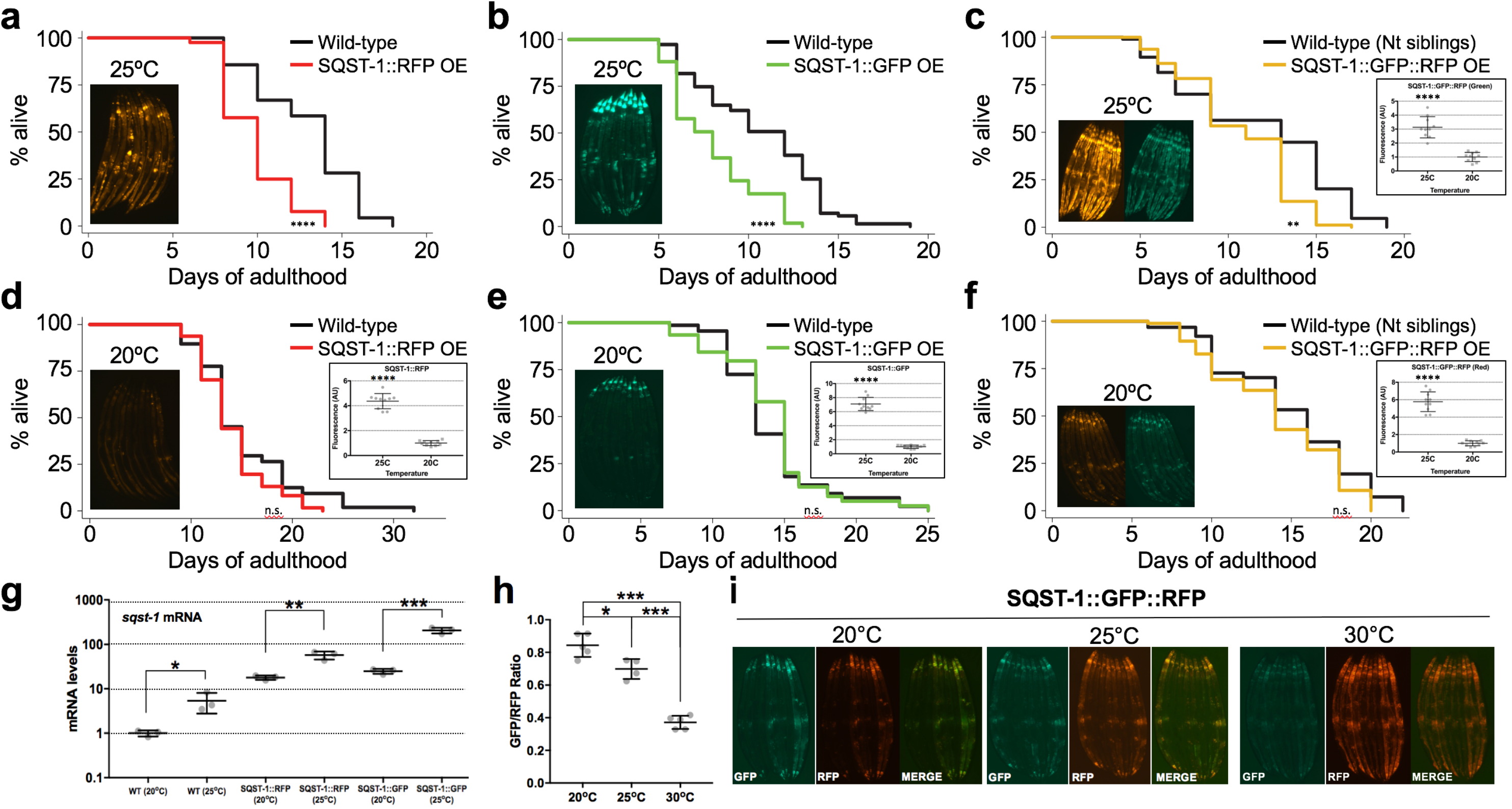
SQST-1 dynamics and lifespan modulation is temperature-dependent in *C. elegans*. **a-f**. Lifespan analysis of wild-type animals and animals over-expressing SQST-1 fused to RFP, GFP or GFP::RFP fed OP50 *E. coli*, developmentally raised at 20°C and then grown at 25°C (**a-c**) or 20°C (**d-f**) during adulthood (n=100). Insets include corresponding representative images of transgenic animals at Day 5 of adulthood, and quantified fluorescence, ±SD *t-test ****p<0.001*. Nt: Non transgenic. **g**. *sqst-1* mRNA levels in wild-type and transgenic animals were quantified by qPCR. Biological triplicates ±SD *t-test *p<0.05, **p<0.01, ***p<0.001* **h**. Levels of GFP and RFP were measured in transgenic tandem SQST-1::GFP::RFP animals (leveraging the pH sensitivity of GFP to visualize SQST-1 accumulation in autolysosome) after incubating Day 1 animals at 20°C, 25°C or 30°C for 24 hours. Average of 10 worms per image. n=5 images per condition ±SD *ANOVA *p<0.05, **p<0.01, ***p<0.001*. Difference in signal between GFP and RFP is due to the low pH sensitivity of GFP. Thus, RFP-only signal represents SQST-1 protein in autolysosomes. **i**. Representative images of transgenic animals (quantified in h.) incubated at 20°C, 25°C or 30°C for 24 hours. Details about lifespan analyses and repeats are available in Supplemental Table 4, Mantel-Cox log-rank. n.s.: not significant, *\*\*p<0.01,****p<0.001*.

### Proteins that bind lipid droplets modulate SQST-1 dynamics

In order to better understand how SQST-1 is regulated, we opted for an unbiased genome-wide RNAi screening approach in an integrated SQST-1 over-expressing strain (*psqst-1::sqst-1::gfp*) at 25°C during development. SQST-1 modulators were subsequently validated by gene silencing in a separate over-expressing strain (*psqst-1::sqst-1::rfp*) during development and adulthood. Interestingly, the vast majority of modifiers led to the intestinal accumulation of fluorescently tagged SQST-1 in both development and adulthood; no modifiers were identified that consistently caused decreased levels of SQST-1 (**Supplemental Table 1**). A substantial portion of genetic modifiers clustered in ribosomal-related proteins, including proteins coding for small and large subunits, suggesting that ribosomal assembly and function substantially impact SQST-1 dynamics (**Figure 2a, Supplemental Table 1**). This is in line with studies in *C. elegans*^2^ and killifish^39^, highlighting the age-related instability of several ribosomal proteins and consequent ribosomal mis-assembly. Loss of ribosome subunit RPL-43 was also found to increase SQST-1 accumulation during development^40^. Since SQSTM1 was also recently reported to associated with lipid droplets in macrophages^41^, we wondered if SQST-1 might also associate with lipid droplets. We co-expressed SQST-1::RFP and the lipid resident protein DHS-3 fused to GFP ^42^ in nematodes and we found that close to 50% of SQST-1::RFP localizes to lipid droplets when autophagy is inhibited (**Figure 2b and Supplemental Figure 2a**). Thus, we investigated whether SQST-1 modulators may also be related to lipid droplet metabolism. We compared our SQST-1 modulators to proteins that have been found to bind lipid droplets^43^ and found significant overlap (*p* < 5.745e-60) (**Figure 2c**). Interestingly, several proteins prone to age-dependent aggregation^2^ also modulated SQST-1 accumulation (*p* < 6.030e-10) (**Figure 2c**). Overall, 17 SQST-1 modulators have the propensity to both aggregate with age and bind lipid droplets (**Figure 2c, Supplemental Table 2)**. Several ribosomal subunits (*rps* and *rpl* genes) and mRNA-related proteins (*inf-1* and *pab-1*) emerged, along with the HSP70 chaperone family member *hsp-1/HSPA8*, indicating that ribosomal assembly, translation initiation, and protein folding are important regulators of SQST-1 dynamics (**Figure 2d**). SQST-1::RFP accumulation upon silencing of these modifiers was mostly gonadal and intestinal (**Figure 2d, Supplemental Figure 2b**). Interestingly, the expression of *sqst-1* was differentially regulated by the 17 SQST-1 modulators whereas silencing large ribosomal proteins generally increased *sqst-1* expression (**Supplemental Figure 2b**). This highlights that *sqst-1* expression is potentially sensitive to ribosomal protein stoichiometry. Here, our comparative analysis suggests that age-related SQST-1 accumulation may be partly attributed to altered interactions between lipid droplets and progressively unstable protein complexes like the ribosome.

**Figure 2.**
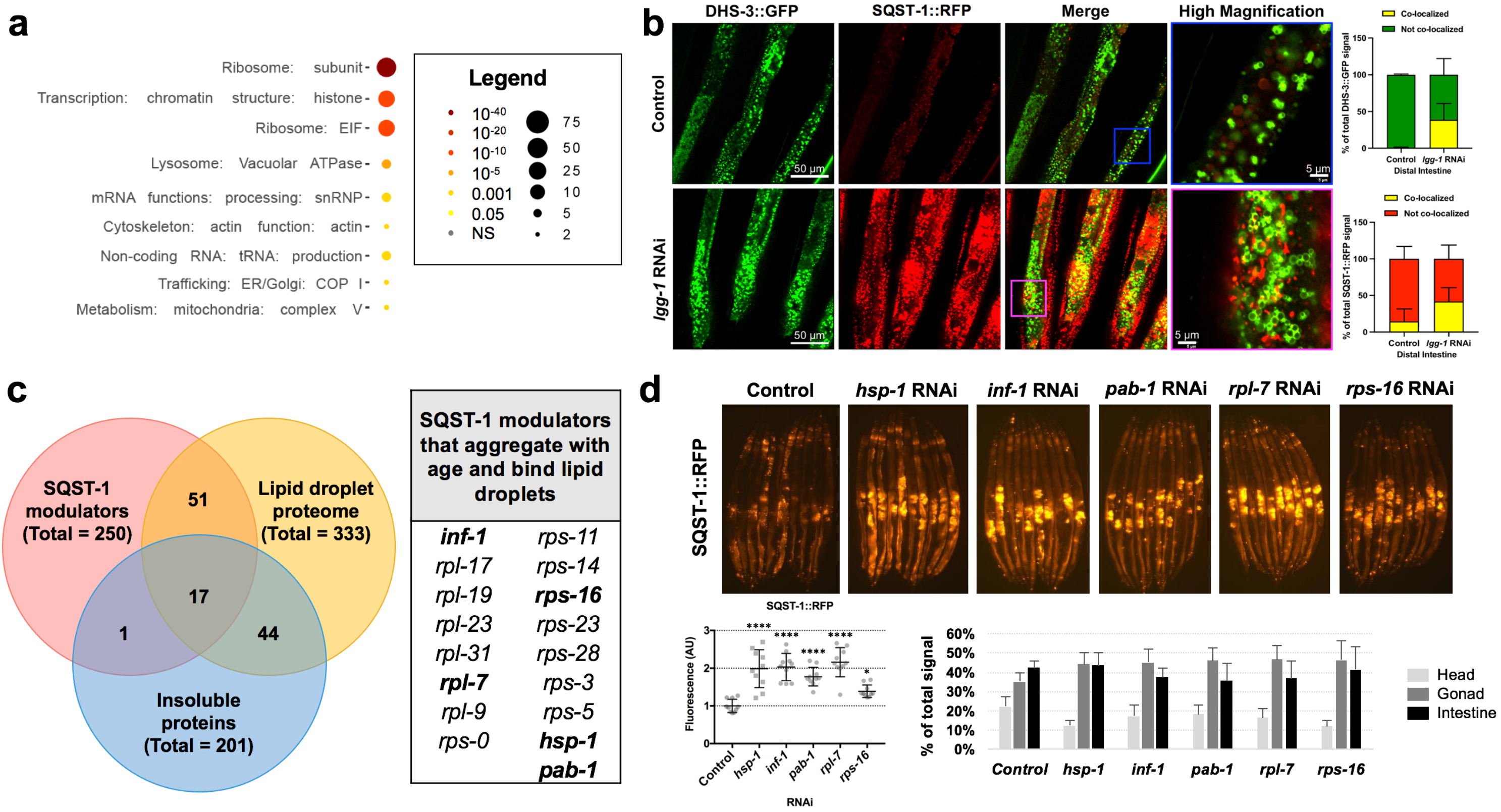
SQST-1 regulators include proteins that bind lipid droplets and that aggregate with age. **a**. Pathway enrichment of genes that regulate SQST-1 levels using WormCat^77^. **b**. SQST-1::RFP accumulates after autophagy is inhibited by *lgg-1* silencing for 3 days during adulthood. Representative confocal microscopy image of the distal intestine of animals expressing both SQST-1::RFP and the lipid droplet-resident protein DHS-3 fused to GFP, subjected to *lgg-1* silencing for 3 days during adulthood. The co-localization between DHS-3::GFP and SQST-1::GFP is quantified. **c**. Overlap of SQST-1 regulators with age-dependent aggregating proteins^2^ and lipid droplet-associated proteins^40^. **d**. Representative images of gene silencing of a set of overlapping SQST-1 regulators during adulthood for 3 days in animals over-expressing SQST-1::RFP. Fluorescence and tissue distribution quantified ±SD *ANOVA *p<0.05,****p<0.001*. Details about SQST-1 regulators are available in Supplemental Tables 1 and 2.

### Elevated lipid droplets reduce SQST-1 accumulation and extend lifespan

Intestinal lipid droplet accumulation is a striking feature of several long-lived nematodes^30-32^. Since SQST-1 accumulated on lipid droplets when autophagy was inhibited (**Figure 2b**), and several SQST-1 modifiers bind lipid droplets (**Figure 2c**), we speculated that lipid droplets may underlie the ability of long-lived animals to stabilize or facilitate clearance of potentially aggregating proteins and maintain proteome stability during aging. In order to specifically stimulate intestinal lipid droplet accumulation and expansion, we silenced the conserved cytosolic lipase *atgl-1/ATGL*^44^ during adulthood to attenuate lipid droplet breakdown, which led to larger lipid stores thereby mimicking a key feature of several long-lived animals (**Figure 3a, inset**). Silencing *atgl-1* resulted in a significant lifespan extension in wild-type animals (12-28%), indicating that lipid droplet accumulation is sufficient to mediate longevity (**Figure 3a-b**). Silencing ATGL-1 activator *lid-1* or hormone-sensitive lipase orthologue *hosl-1*^45^ also led to significant reduction in SQST-1 accumulation (**Supplemental Figure 2c**). These observations pointed to a possible lipid droplet-mediated cytoprotective mechanism^46^. Indeed, silencing *atgl-1* in SQST-1::RFP over-expressing animals led to a substantial increase in lifespan (**Figure 3c)** accompanied by a marked decrease in SQST-1 accumulation (**Figure 3d**). Similar observations were made in animals over-expressing SQST-1::GFP (**Supplemental Figure 2d**), suggesting that the lipid droplet-mediated effects impacts SQST-1-mediated selective autophagy.

**Figure 3.**
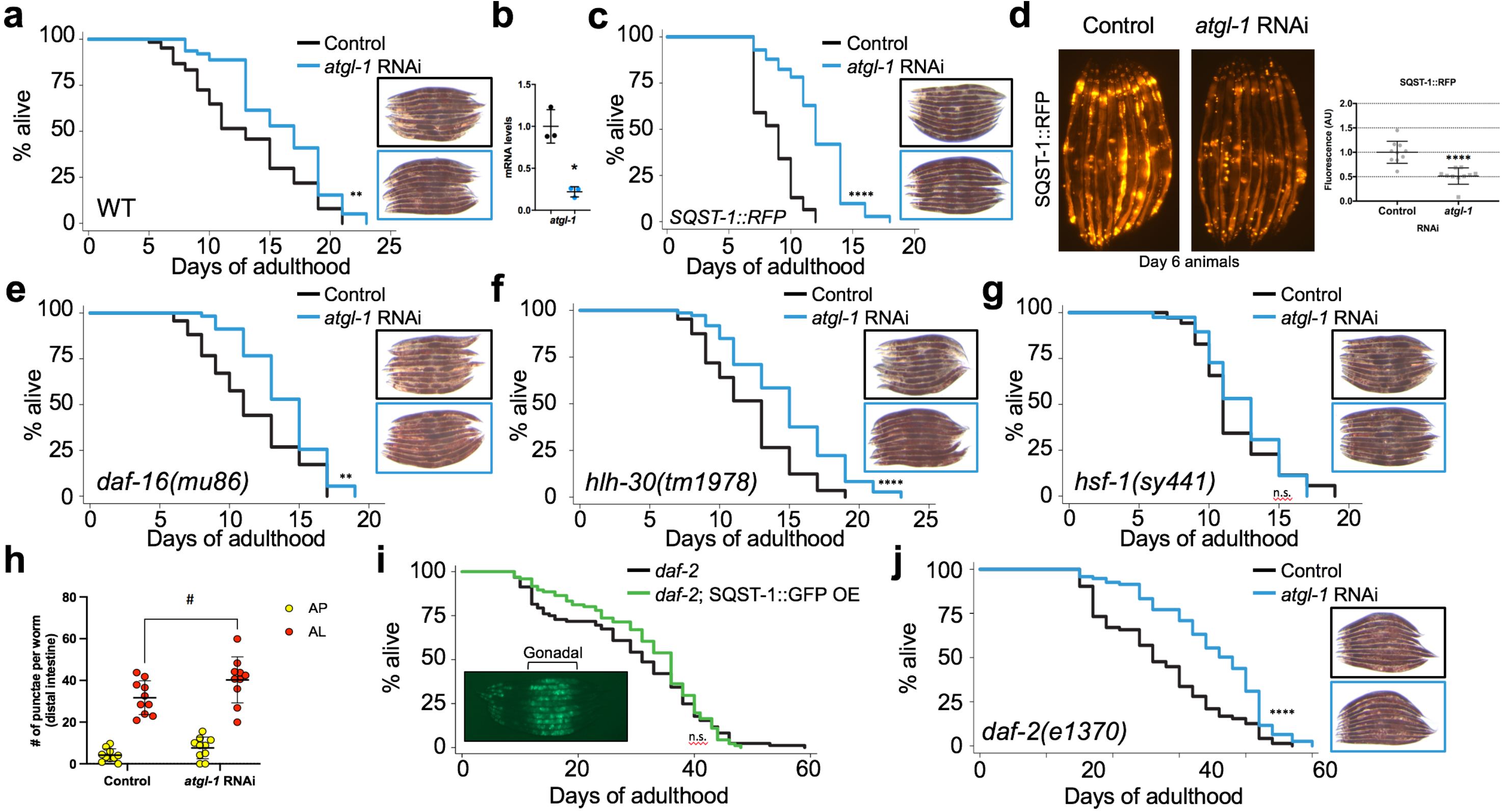
Silencing *atgl-1* extends lifespan and reduces SQST-1 accumulation in *C. elegans*. **a**. Lifespan analysis of wild-type animals and **c**. animals over-expressing SQST-1::RFP raised at 20°C on OP50 *E. coli* and then grown at 25°C during adulthood on control bacteria or bacteria expressing dsRNA against *atgl-1* (n=100). Corresponding images of Day 5 animals stained with Oil-Red-O demonstrating intestinal lipid store accretion under *atgl-1* RNAi (blue borders). **b**. Levels of *atgl-1* mRNA measured by qPCR showing efficient silencing after 3 days on *atgl-1* RNAi (Black: Control RNAi, Blue: *atgl-1* RNAi). Biological triplicates *t-*test *\*p<0.05* **d**. Representative fluorescence images taken after Day 1 animals over-expressing SQST-1::RFP and DHS-3::GFP were fed control bacteria or bacteria expressing dsRNA against *atgl-1* for 5 days at 25°C. **e-g**. Lifespan analysis of *daf-16(mu86), hlh-30(tm1978)*, and *hsf-1(sy441)* mutants raised at 20°C on OP50 *E. coli* and then grown at 25°C during adulthood on control bacteria or bacteria expressing dsRNA against *atgl-1* (n=100). Corresponding images of Day 5 animals stained with ORO demonstrating intestinal lipid store accretion under *atgl-1* RNAi (blue borders). **h**. Levels of autophagosomes and autolysosomes were measured in animals expressing the tandem reporter mCherry::GFP::LGG-1^7^ after feeding Day 1 animals with control bacteria or bacteria expressing dsRNA against *atgl-1* for 2 days at 25°C. n=10 per condition ±SD *t-*test ^*#*^*p<0.06* **i**. Lifespan analysis of *daf-2(e1370)* and *daf-2;SQST-1::GFP* animals raised at 20°C and then grown at 25°C during adulthood on OP50 *E. coli* (with representative image of Day 5 animals, comparative image of wild-type animals in Figure 1b). **j**. *daf-2(e1370)* animals were raised at 20°C on OP50 *E. coli* and then grown at 25°C during adulthood on control bacteria or bacteria expressing dsRNA against *atgl-1* (n=100). Corresponding images of Day 5 animals stained with ORO demonstrating intestinal lipid store accretion under *atgl-1* RNAi (blue borders). Details about lifespan analyses and repeats are available in Supplemental Tables 3 and 4, Mantel-Cox log-rank. n.s.: not significant, *\*\*p<0.01,****p<0.001*..

To determine whether accumulation of lipids is able to improve the lifespan of proteostatically-impaired mutants, we silenced *atgl-1* in three well-established short-lived mutants, including *daf-16(mu86), hlh-30(tm1978)*, and *hsf-1(sy441)*^47^. Notably, the expression of these transcription factors was increased at 25°C compared to 20°C, suggesting that their function may be stimulated by heat (**Supplemental Figure 2e**). Silencing *atgl-1* extended the lifespan of *daf-16* and *hlh-30* mutants, but not the lifespan of *hsf-1* mutants (**Figure 3e-g**), indicating that lipid droplet-mediated lifespan extension may require the expression of key chaperones regulated by HSF-1/HSF1, such as heat shock protein HSP-1, which binds lipid droplets and modulates SQST-1 levels (**Figure 2d**). Interestingly, lipid droplets may harbor chaperones with roles in aggregate clearance^48^. Notably, autophagic activity, as measured by the tandem mCherry::GFP::LGG-1 reporter^7^, showed that *atgl-1* silencing increases the conversion of autophagosomes (GFP+mCherry - yellow puncta) into autolysosomes (mCherry only - red puncta) (**Figure 3h, Supplemental Figure 2f**). Accordingly, the lifespan of autophagy-deficient *atg-7* mutants was not increased by silencing *atgl-1* (**Supplemental Figure 2g**).

As our lifespan analyses highlighted novel proteostatic and longevity roles for lipid droplets, we reasoned that long-lived animals with large lipid stores should accumulate less intestinal SQST-1. Strikingly, *daf-2(e1370)* expressing SQST-1::GFP (**Figure 3i**) or SQST-1::RFP (**Supplemental Figure 2h**) accumulated negligible intestinal SQST-1 during aging (GFP signal in mid-section was entirely gonadal, see wild-type comparison in **Figure 1b**). In addition, SQST-1 over-expression did not significantly affect the long lifespan of *daf-2* animals (**Figure 3i**). Loss of *sqst-1* did not affect the lifespan of *daf-2* animals (**Supplemental Figure 2i**), as previously shown^16^, highlighting that SQST-1 function becomes less important at 25°C for the lifespan of organisms with relatively stable proteomes and high lipid stores. The extent of heat-induced increase in SQST-1-mediated selective autophagy was also attenuated in *daf-2* animals (**Supplemental Figure 2j**) compared to wild-type (**Figure 1h**), suggesting that elevated lipid droplets may buffer the need for SQST-1 function during heat stress. Silencing *atgl-1* in *daf-2* animals enhanced their intestinal lipid stores and further extended their lifespan (**Figure 3j**), indicating that elevated lipid droplet accumulation can also extend lifespan in animals with enhanced proteostasis. Altogether, our data present an important and previously unrecognized role for lipid droplets in SQST-1 dynamics and longevity.

### Silencing atgl-1 elicits limited changes in gene expression

Lipid droplets have been recently shown to coordinate transcriptional programs by sequestering factors that modulate transcription^49^. Since HSF-1 was required for lifespan extension by *atgl-1* silencing, we hypothesized that the lipid droplet increase associated with *atgl-1* silencing might affect the expression of HSF-1-regulated targets. Using RNA sequencing (RNA-seq), we found that enhancing lipid stores by silencing *atgl-1* in WT or *daf-2* animals had limited effect on global transcription. Silencing *atgl-1* resulted in 13 overlapping differentially expressed genes (DEG) in wild-type animals and *daf-2* mutants of which 7 genes are down-regulated in both cases (**Supplemental Figure 3a-b**). The mRNA levels of *sqst-1* remained unchanged by *atgl-1* silencing (**Supplemental Figure 3c**). Altering *atgl-1* levels (silencing or over-expression) led to differential expression of 50 overlapping genes, with no discernable transcription factor signature (**Supplemental Figure 3d-e**). In addition, expression of *sqst-1* was unchanged in wild-type animals with low or high levels of *atgl-1* (**Supplemental Figure 3e**). Overall, we concluded that transcriptional regulation may not contribute significantly to the proteostatic-enhancing and lifespan-extending effects of lipid droplets. Therefore, lipid droplets may impact proteostasis via a more direct mechanism on the proteome itself.

### Lipid droplets enhance proteostasis by modulating the accumulation of ubiquitinated proteins

The emerging connection between lipid droplet stores and proteome stability led us to investigate how lipid droplet loss affects proteostasis and SQST-1 abundance. First, we tested whether lipid droplet depletion affects lifespan using ATGL-1::GFP over-expressing animals^50^. Unlike at 20°C (**Supplemental Table 3**)^51^, we found that over-expressing ATGL-1 was detrimental to lifespan at 25°C (**Figure 4a**) and led to a significant increase in SQST-1 intestinal accumulation accompanied by lower lipid stores (**Figure 4b**), suggesting that loss of lipid droplet stores can interfere with SQST-1 dynamics. When autophagy or proteasome function was reduced by silencing autophagosome protein *lgg-1* or proteasome subunit *rpn-6.1*, respectively, ubiquitinated protein levels were higher in animals with lower lipid stores, suggesting that lipid droplets may buffer the proteome and facilitate the processing of unstable or misfolded proteins (**Figure 4c**). Strikingly, analyzing lipid droplets revealed preferential accumulation of ubiquitinated proteins in lipid droplet-enriched fractions (**Figure 4d, Supplemental Figure 4a**), indicating that lipid droplets contain a significant amount of proteins bound for degradation. Lipid droplet-associated accumulation of ubiquitinated proteins was increased when autophagic or proteasomal degradation was reduced (**Figure 4d**), suggesting that lipid droplets have the capacity to harbor many degradation cargoes, which may become particularly relevant for proteostasis when autophagic and proteasomal systems are failing during aging.

**Figure 4.**
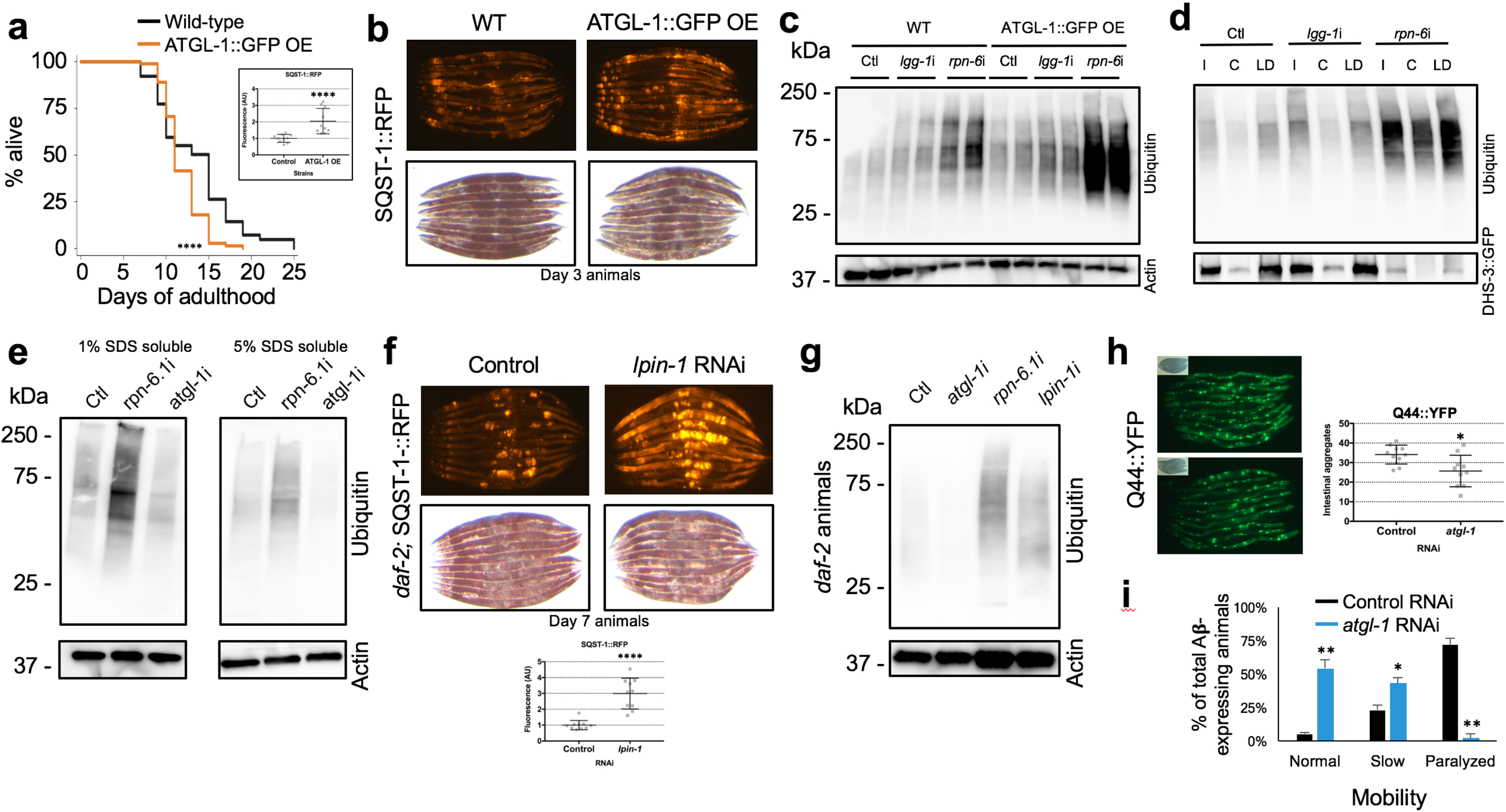
Lipid droplets modulate protein ubiquitination levels and enhance proteostasis. **a**. Lifespan analysis of wild-type and transgenic animals over-expressing ATGL-1::GFP developmentally raised at 20°C and then grown at 25°C during adulthood on OP50 *E. coli* (n=100). **b**. Levels of RFP signal and lipid droplets from Day 3 animals over-expressing SQST-1::RFP in wild-type or transgenic ATGL-1::GFP over-expressing background. **c**. Wild-type and ATGL-1::GFP over-expressing animals were raised at 20°C and then grown at 25°C during adulthood on control bacteria (Ctl) or bacteria expressing dsRNA against autophagy gene *lgg-1* or proteasome subunit gene *rpn-6.1* for 2 days. Levels of ubiquitinated proteins and actin were visualized by immunAnimals expressing lipid droplet-resident protein DHS-3 fused to GFP were raised at 20°C and then grown at 25°C during adulthood on control bacteria or bacteria expressing RNAi against *lgg-1* or *rpn-6* for 4 days. Levels of ubiquitinated proteins and DHS-3::GFP were visualized by immunoblotting from total input (I), cytosol (C) and lipid droplet (LD) fractions (comparative % loaded between fractions, i.e. 10%). oblotting. Biological replicates shown. **d. e**. Day 1 wild-type animals were fed control bacteria or bacteria expressing dsRNA against *rpn-6* or *atgl-1* for 3 days and ubiquitination levels were detected by immunoblotting. **f**. Levels of RFP signal and lipid droplets in Day 7 *daf-2* animals over-expressing SQST-1::RFP and fed control bacteria or bacteria expressing dsRNA against *lpin-1* during adulthood. **g**. Day 1 *daf-2* animals were fed control bacteria or bacteria expressing dsRNA against *atgl-1, rpn-6* or *lpin-1* for 5 days at 25°C and ubiquitination levels were detected by immunoblotting. **h**. Nematodes expressing intestinally Q44::YFP were grown at at 20°C on OP50 *E. coli* and fed control bacteria or bacteria expressing dsRNA against *atgl-1* at Day 1 of adulthood for 4 days at 25°C, followed by quantification of intestinal aggregates. ±SD *t-*test *\*p<0.05*. **i**. Nematodes expressing heat-inducible human Aβ-42 were grown at 20°C on OP50 *E. coli* and fed control bacteria or bacteria expressing dsRNA against *atgl-1* at Day 1 of adulthood for 2 days at 25°C. Paralysis was scored thereafter. Triplicates of n=100 each, ±SD *t-*test *\*p<0.05, **p<0.01*.

The association and function of ATGL-1 with lipid droplets was recently found to be antagonized by the AAA-ATPase CDC-48/VCP^52^. CDC-48/VCP is best known for its role in ER-associated degradation machinery^53^, but it has been associated with other functions including endocytosis^54^ and more recently autophagy itself^55^. CDC-48/VCP helps unfold unstable, ubiquitinated, and proteasomal degradation-bound proteins in concert with heat shock protein HSP-70, a key regulated target of HSF-1^56^. Thus, we assayed the effect of the loss of CDC-48 on SQST-1 and ATGL-1 levels using fluorescent reporters. Silencing *cdc-48.2* increased the levels of ATGL-1::GFP (**Supplemental Figure 4b)** and led to the accumulation of SQST-1::RFP (**Supplemental Figure 4c**). In mammalian cells, loss of proteasome function drives SQSTM1 to divert cargo to selective autophagy^57^, but stimulating proteasomal processing by over-expressing of VCP can reduce SQSTM1 accumulation^58^. Here, we reasoned that CDC-48 functions related to processing ubiquitinated targets and autophagy may underlie the ability of nematodes with elevated lipid droplets to attenuate SQST-1 accumulation and modulate lifespan. Accordingly, silencing *atgl-1* in *cdc-48.1* or *cdc-48.2* mutants failed to significantly extend lifespan (**Supplemental Figure 4d, Supplemental Table 4**), indicating that lipid droplet-mediated lifespan extension requires functional CDC-48.

As lipid droplet size increases, its surface also increases, and it is possible that the capacity of this organelle to bind and stabilize proteins may also increase as well. Enhancing lipid droplet stores by silencing *atgl-1* reduced the overall accumulation of ubiquitinated proteins, in particular in the lower solubility (5% SDS soluble) fraction (**Figure 4e**), suggesting that typically insoluble proteins are less likely to be ubiquitinated when lipid droplets abound. Notably, the types of proteins that aggregate with age differ between wild-type and long-lived *daf-2* mutants, as the latter tends to accumulate aggregating proteins that are less hydrophobic than those aggregating in wild-type animals^4^. Reducing lipid droplet stores genetically by silencing lipogenic genes *sbp-1(SREBP2), lpin-1(LIPN1)* or *fasn-1(FASN)* enhanced SQST-1 accumulation in wild-type animals (**Supplemental Figure 4e**). Similarly, loss of lipid droplet stores by silencing *lpin-1* in *daf-2* resulted in increased SQST-1 accumulation and overall protein ubiquitination (**Figure 4f-g**). As proteostasis failure is a feature of neurodegeneration, we tested the effect of increasing lipid droplet levels in proteotoxic contexts. Animals intestinally-expressing an aggregating poly-glutamine protein fused to YFP (Q44::YFP)^59^ had reduced numbers of aggregates when *atgl-1* was silenced **(Figure 4h)**. Reducing the expression of *atgl-1* in a muscle-expressing proteotoxic amyloid protein Aβ-42 resulted in a marked protection against aggregation-associated paralysis^60^ (**Figure 4i**). Altogether, our data demonstrate that lipid droplets are important for proteostasis and contribute to lifespan by facilitating autophagy and stabilizing the proteome.

## DISCUSSION

Decades of aging research has uncovered key longevity-regulating pathways in *C. elegans*^47^, yet the role of lipid droplets in the lifespan of several established long-lived nematodes has remained unresolved. Here, while studying the regulation of the selective autophagy receptor SQST-1, we unexpectedly uncovered a direct role for lipid droplets in proteostasis and lifespan. Several SQST-1 modulators associate with lipid droplets and aggregate with age, including ribosomal, translation-related, and folding-related proteins^43^. Ribosomal assembly dysfunction due to loss in subunit stoichiometry may burden the autophagy machinery, and contribute to age-related proteotoxicity^39^. Importantly, lipid droplet accumulation, by silencing cytosolic triacylglycerol lipase *atgl-1/ATGL*, mitigates the progressive age-related SQST-1 accumulation, a benefit recapitulated in long-lived, lipid droplet-rich *daf-2* mutant animals. Lipid droplets are cytoprotective as they prevent aberrant SQST-1 accumulation, in part via the functioning of AAA-ATPase CDC-48/VCP which processes ubiquitinated proteins for proteasomal degradation^61^. Strikingly, we found that the majority of soluble ubiquitinated proteins in nematodes was associated with lipid droplets, suggesting that unstable proteins bound for degradation may be stabilized by the surface of lipid droplets. Our findings also provide support to the importance of the emerging lipid-droplet-mediated protein degradation process^62,63^ in longevity and aging. Altogether, our study strengthens the emerging concept that lipid droplets serve as a buffer for proteostasis^46^ by stabilizing proteomes and coordinating protein degradation machineries.

Understanding the regulation of SQST-1/SQSTM1 during aging and in different disease contexts is important in order to determine the validity of stimulating selective autophagy as a therapeutic strategy to improve proteostasis in age-related diseases^64^. Here, we find that over-expressing SQST-1 had no effect on the lifespan of wild-type animals at 20°C^16^ whereas elevated SQST-1 level at 25°C was detrimental. In hindsight, one may have predicted that over-expressing a ubiquitin-binding protein with a propensity for oligomerization^65^ might not be necessarily advantageous, particularly in organisms nearing their protein solubility limit^2,3^ and displaying overall proteome instability during aging^5^. SQST-1 over-expression was primarily visible in neuronal, gonadal and intestinal cells and accumulated in a temperature-dependent manner and progressively during aging. The inability of stressed and aging animals to process ubiquitinated cargos bound for proteasomal or autophagic degradation may lead to a gradual build-up of SQST-1. Accordingly, SQST-1 over-expression may over-sensitize the animals to unstable, ubiquitination-prone proteins, posing an additional challenge to the autophagic machinery. This is exemplified by recent evidence that reducing SQST-1 specifically in the neurons of a nematode model of ALS is protective^66^, possibly since mutant fused in sarcoma (FUS) expression promotes SQST-1 accumulation, which exacerbates proteotoxicity and neurodegeneration. Notably, SQSTM1 can phase separate when interacting with ubiquitinated cargoes^67^, but the impact of the formation of these condensates on overall proteostasis is unclear.

Lipid droplet accumulation is a prominent and under-explored phenotype of several long-lived nematodes, including well-established models such as germline-less *glp-1* animals, protein translation *rsks-1* mutants, insulin/IGF-1 receptor *daf-2* mutants^68^ as well as in long-lived animals with reduced nuclear export^69^. In addition, lifespan extension by inhibiting intestinal lipid secretion mediated by vitellogenins results in lipid redistribution to intestinal lipid droplet stores^29^, a process believed to generate precursors for lysosomally-derived lipid signals^28^. While enhanced lysosomal lipolysis and lipophagy may stimulate longevity-associated lipid signaling, overactive cytosolic lipolysis decreases the ability of cells to maintain a stable proteome, which may burden the selective autophagy machinery. Our findings support that lipid droplets are much more than just lipid storage organelles^46^ and can serve as interfaces to enable protein stabilization and processing.

Rationing of intestinal lipid stores was originally evoked as a potential mechanism to provide long-term energy for dauer larvae that arrest feeding^35^. Here, we propose a role for lipid stores during adulthood that focuses on their capacity to buffer proteostasis during aging in concert with heat shock chaperones, thereby unburdening protein degradation systems (**Figure 5**). ATGL-1 over-expression leads to lipid store depletion ^51^, but, more importantly, it also removes the protection conferred by lipid droplets and renders animals vulnerable to proteotoxic stress associated with reduced proteasomal function, heat and aging. Thus, regulating the activity of ATGL-1 via transcription^70^, post-translational modification^71^ or alteration in CDC48/VCP activity^52^ provide a mechanism by which cells modify protein homeostasis. Overall, our data also show that the lipid droplet-mediated decrease in protein ubiquitination and the enhancement in lifespan require the function of the AAA-ATPase CDC-48/VCP, a ubiquitin protein processing enzyme and autophagy modulator^55^.

**Figure 5.**
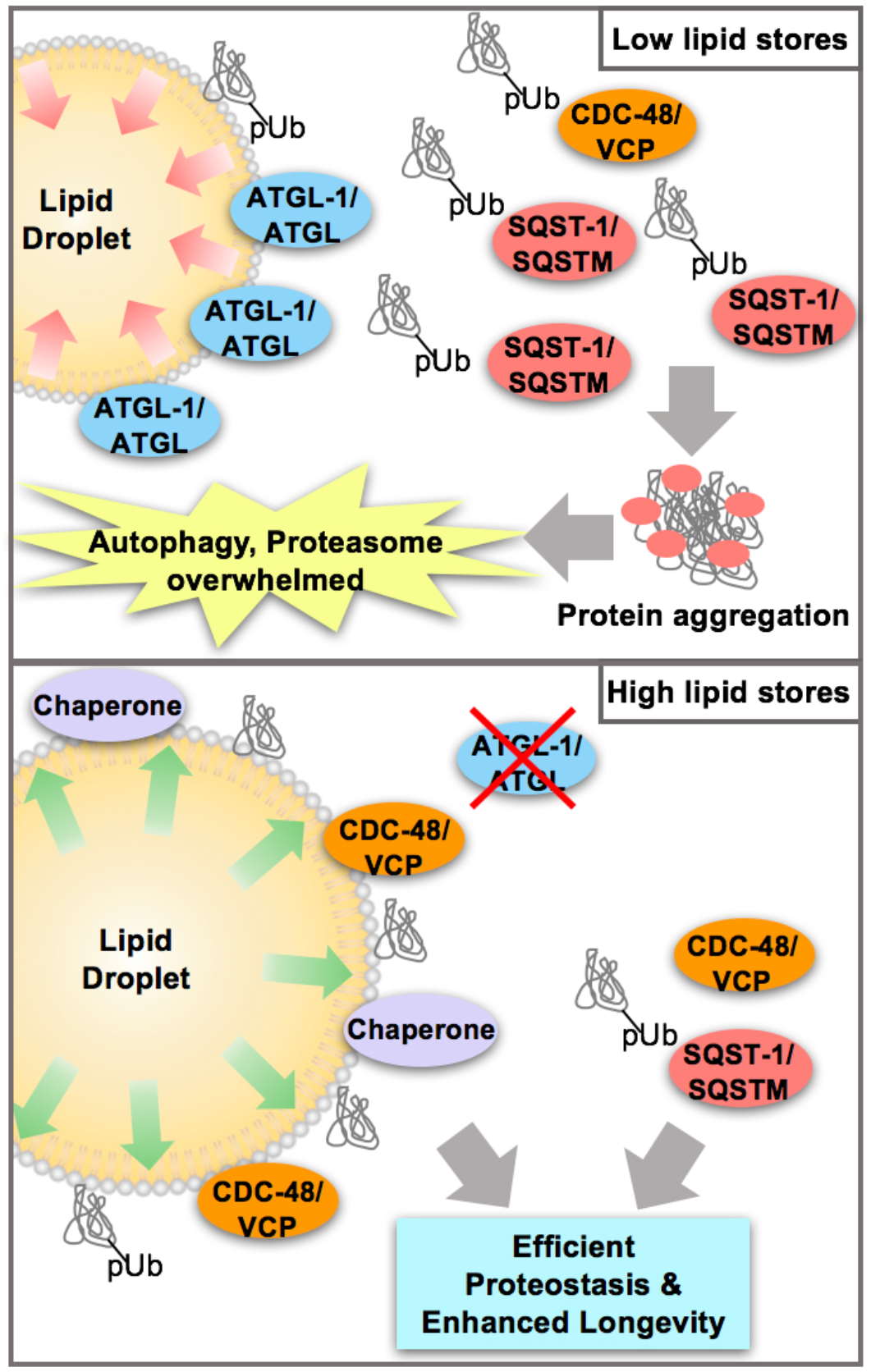
Lipid droplets buffer proteostasis by harboring proteins that challenge protein degradation pathways. When lipid stores are low, ubiquitinated proteins accumulate and eventually overwhelm the proteasomal system and selective autophagy receptors such as SQST-1. High lipid droplet content increases the capacity of cells to stabilize proteins (in conjunction with CDC-48 and HSF-1-regulated chaperones, such as HSP-1) that would otherwise accumulate and aggregate. As proteostasis declines with age, lipid droplets can facilitate autophagic degradation and stabilize the proteome, which prevents the burdening of selective autophagy.

Lipid droplet expansion may belong to an arsenal of proteostatic measures that lipid-storing cells and possibly neurons can employ to prevent protein misfolding and aggregation. While lipid droplet accumulation in immune cells such as macrophages^72^ and microglia^73^ may be inflammatory and contribute to aging, lipid droplets may confer some post-mitotic, differentiated cells with added protection against protein aggregation^48^ and mitochondrial lipid overload^74^. For example, lipid droplet accumulation in Alzheimer’s disease pathogenesis^75^ may highlight an effort from neuronal cells to improve proteome stability and resilience against progressive proteotoxicity. Notably, drastic inhibition of triacylglycerol lipolysis becomes detrimental to neurons^76^ as larger lipid droplets are less easily broken down^50^. However, a recent study showed that lipid droplet accumulation protected hyper-activated neurons from cell death^77^, a phenomenon potentially relevant in Alzheimer’s disease^78^. Thus, moderate accumulation of lipid droplets may be more desirable to provide protection against age-related proteotoxicity. In agreement, our data show marked protection against paralysis in a proteostatic Alzheimer’s disease model by attenuating cytosolic lipolysis.

Enhancing proteostasis by stimulating the autophagy process has emerged as an attractive strategy to mitigate several age-related diseases with pathological proteostatic decline such as neurodegenerative diseases ^79^. However, our study indicates that stimulating selective autophagy by specifically increasing the expression of one selective autophagy receptor, SQST-1/SQSTM1, is not sufficient to improve proteostasis and may exacerbate age-related proteostatic collapse. Overall, our study suggests that therapeutic improvement in proteostasis may benefit from a combinatorial approach in which the whole autophagy/lysosomal machinery is stimulated concomitantly with a modest reduction in lipid droplet breakdown. Such an approach may prevent pathogenic burdening of the autophagy/lysosome pathway with cargoes that could have otherwise been stabilized by lipid droplets or routed to the proteasome.

## METHODS

### *C. elegans* strain maintenance

Nematodes were maintained at 20°C on agar NGM plates seeded with OP50 *E. coli* unless otherwise noted, as previously described^80^. Synchronized populations were prepared using a sodium hypochlorite solution to collect eggs as previously described^81^. Supplemental Tables 5 and 6 contains the list of strains used in this study. HT115 *E. coli* and RNAi clones from the Ahringer library (Source Bioscience) were used for RNAi experiments^82^.

### Transgenic strain construction

DNA constructs for the plasmids pLAP26 (*psqst-1::sqst-1::rfp::unc-54 3’UTR*) and pLAP29 (*psqst-1::sqst-1::gfp::rfp::unc-54 3’UTR*) were assembled using HiFi cloning and were injected into the germline of Day 1 adults (pLAP26 was co-injected with the pLAP7 (*myo-2::gfp::unc-54)*). Transgenic progeny was UV-irradiated for extrachromosomal array integration and selected for 100% transmission rate and backcrossed at least four times to wildtype N2. Details about strain construction are provided in Supplemental Table 6.

### Genome-wide RNAi screen

Approximately 50 synchronized transgenic eggs (*psqst-1::SQST-1::GFP*) were transferred onto plates seeded with RNAi against all genes from the Ahringer library (Source BioScience), which represents about 86% of the predicted genes in the *C. elegans* genome^82^. Nematodes were developed at 25°C and changes in GFP intensity or expression pattern were monitored on Day 1 of adulthood. As a negative control, bacteria expressing the L4440 backbone (i.e. empty vector) expressed in HT115 *E. coli* were used. The identity of each RNAi was not known to the scorer and fluorescence intensity was scored using a V20 dissection fluorescence microscope (Zeiss). Modifiers from the genome-wide RNAi screen were sequenced (GENEWIZ), validated three times and corroborated using transgenic animals expressing *SQST-1::RFP*. RNAi against the genes with the strongest effect on GFP expression were subsequently validated by silencing in adulthood only. Details are provided in Supplemental Tables 1 and 2.

### Lifespan analyses

Eggs obtained from bleaching were transferred onto OP50 *E. coli*. bacteria-seeded agar plates. All lifespan analyses were carried out at 25°C starting at Day 1 of adulthood after development at 20°C, unless otherwise noted. For gene knockdown experiments, worms were transferred to control bacteria expressing the empty vector (L4440) or expressing corresponding RNAi at day one of adulthood. Viability was scored every 1 to 3 days, as previously described^83^. Survival curves and statistical analysis were generated using the Stata 15.0 software (StataCorp). Details are provided in Supplemental Tables 3 and 4.

### Comparative analyses

Knocked-down genes with the strongest effect on SQST-1::GFP levels were analyzed for gene ontology using WormCat^84^ to identify which pathways were enriched. RNAi against genes that overlapped with the SQST-1 modulators, lipid droplet proteome^43^, and insoluble proteins^2^ were then tested in adult-only experiments. Statistical significance of the overlap between any two groups of genes was calculated by hypergeometric probability (nemates.org, Lund laboratory, University of Kentucky). For adult-only RNAi validation, eggs from wild-type and SQST-1 transgenic animals were developed on OP50 *E coli*. bacteria at 20°C and transferred to corresponding RNAi plates at Day 1 of adulthood and grown at 25°C for an additional 72 hours. Adults were transferred away from their progeny and bright field and fluorescent images were taken daily.

### Imaging and analyses

Worms were visualized using a Zeiss Discovery V20 fluorescence dissecting microscope (Zeiss, White Plains, NY). Worms were immobilized with 0.1% sodium azide in M9 solution (42 mM Na2HPO4 22 mM KH2PO4 86 mM NaCl, 1 mM MgSO4-7H2O) on agar plates and images were taken via Zen imaging software, using consistent parameters (magnification and exposure) within each experiment. For quantification of fluorescent signal, total fluorescence of each worm in each image was analyzed using ImageJ software and averaged for mean total fluorescence. Confocal microscopy images were obtained on an Olympus FV3000 inverted confocal laser scanning microscope (Leduc Bioimaging Facility, Brown University). Quantification of autophagosomes marked by GFP and mCherry showing yellow puncta and autolysosomes marked with only mCherry as GFP is quenched in acidic pH showing up as red puncta in the LGG-1 tandem reporter strain was carried out by color thresholding composite images for yellow or red puncta followed by counting the thresholded puncta using particle analysis on ImageJ. The overlap between lipid droplet protein (DHS-3) and tagged SQST-1 in the over-expressing strain was also performed using color thresholding followed by particle analysis functions in ImageJ (NIH).

### Lipid staining

Worms were collected and washed twice with M9 solution and then fixed with 60% isopropanol for 30 minutes. After fixation, worms were stained overnight on a rocker with freshly prepared 60% Oil Red O (stock solution of 0.5% ORO in isopropanol diluted with water, equilibrated overnight on a rocker, and gravity-filtered). The following day, worms were washed with TBS-T (50 mM Tris base, 150 mM Tris HCl, 1.5 M NaCl, and 0.05% Tween-20) and imaged using a Zeiss Discovery V20 fluorescence dissecting microscope.

### Gene expression analysis

RNA was extracted from approximately 3,000 Day 1 worms and cDNA was prepared as previously described ^8^. Gene expression levels were measured in biological triplicates using iTaq Universal SYBR Green Supermix (BIO-RAD), and a Roche LightCycler 96 (Indianapolis, IN). Expression was normalized using 2 housekeeping genes, *act-1* and *cyn-1* and statistical analyses were performed using GraphPad Prism 7 (GraphPad Software). See Supplemental Table 7 for qPCR primers details.

### RNAseq sample preparation and analysis

WT or *daf-2* mutant worms were developed on OP50 at 20°C and transferred to corresponding RNAi plates at Day 1 of adulthood and grown at 25°C for an additional 96 hours. Worms over-expressing ATGL-1::GFP were developed on OP50 at 20°C and grown at 25°C for an additional 96 hours. Adults were transferred away from their progeny daily. RNA was extracted and cDNA was prepared as previously described^8^. RNA quality was confirmed by BioAnalyzer (Genomics Core, Brown University), and *atgl-1* silencing was confirmed by qPCR before submitting samples for RNAseq analysis by GENEWIZ as previously described^8^.

### Lipid droplet fractionation

Lipid droplets were isolated by ultracentrifugation from 24,000 Day 5 animals grown at 25°C on RNAi for 96 hours. Worms were homogenized with a ball-bearing homogenizer in MSB buffer (250mM sucrose and 10mM Tris-HCl pH 7.4) containing protease inhibitor (Roche). The lysate was cleared of cuticle debris and unlysed worms by centrifuging at 500 x g for 5 minutes. 10% of the cleared lysate was reserved as “Input”. Nuclei and non-lipid droplet organelles were pelleted out of the cytosol by an ultracentrifugation spin at 45,000 RPM in a TLA100.1 rotor for 30 minutes at 4°C. The lipid droplet layer at the top of the centrifugation tube was aspirated, and contaminating organelles were removed by an additional centrifugation of the lipid droplet fraction. The final volume of lipid droplet and cytosol fractions were kept consistent across conditions.

### Immunoblotting

Protein lysates were collected from worms using RIPA buffer (50 mM Tris-HCl, 250 mM sucrose, 1 mM EDTA, and Roche protease inhibitor tablet, with 1% or 5% SDS) and a handheld homogenizer, and cleared lysate protein concentrations were quantified using the DC Protein Assay kit (BIO-RAD). Equal amounts of total protein (10 μg) or comparative volumes for lipid droplet fractions (Input volume is 10% of lipid droplet and cytosol fractions) were separated by SDS-PAGE on a 4-15% Tris-Glycine gel and transferred to nitrocellulose. The membrane was briefly stained with Ponceau S to confirm even transfer, rinsed with TBS-T until clear, blocked with 5% nonfat dry milk in TBS-T, and immunoblotted with anti-Ubiquitin (Invitrogen MAI-10035), anti-GFP (Santa Cruz SC-8334), and anti-Actin (Milllipore, MAB1501R). The membrane was developed using SuperSignal West Femto Maximum Sensitivity Substrate (ThermoFisher) and imaged on a ChemiDoc Imaging System (BIO-RAD).

## DATA AVAILABILITY STATEMENT

RNA sequencing data are accessible on NCBI GEO (Accession number: GEO204953).

## CONTRIBUTIONS

AVK, JM, WMP and JAL performed lifespan analyses and imaging as well as RNAi sequencing, validation, and analysis. AVK, JM, JAL and JRJ validated the whole-genome RNAi screening. JM performed the RNA sequencing and lipid droplet analyses. WMP conducted lipid staining and AVK performed the confocal microscopy imaging. DIR performed RNAi sequencing and conducted qPCR analyses. CN, RP and JLA performed lifespan analysis repeats. SQW and WMP developed SQST-1 over-expressing and tandem reporter strains. LRL designed the experiments, conducted initial RNAi screening, imaging, qPCR and lifespan analyses, and wrote the manuscript. All co-authors edited the manuscript.

## ACKNOWLEDGEMENTS

We are grateful for the technical support provided by Erin McConnell, Rachel Tam and Tuong Tran. We thank the Malene Hansen laboratory (SBPMDI) and the Caenorhabditis Genetics Center (U. Minnesota, P40 OD010440) for providing transgenic and mutant strains and the Andrew Dillin laboratory (UC Berkeley/HHMI) for sharing expression plasmids. This work was funded by grants from the National Institute of Health (R00 AG042494, R01 AG051810 and R21 AG068922) and a Glenn Award for Research in Biological Mechanisms of Aging from the Glenn Foundation for Medical Research to LRL.

