## Supplemental Figures and Tables for "Lipid droplets modulate proteostasis, SQST-1/SQSTM1 dynamics, and lifespan in *C. elegans*"

Supplemental Figure 1

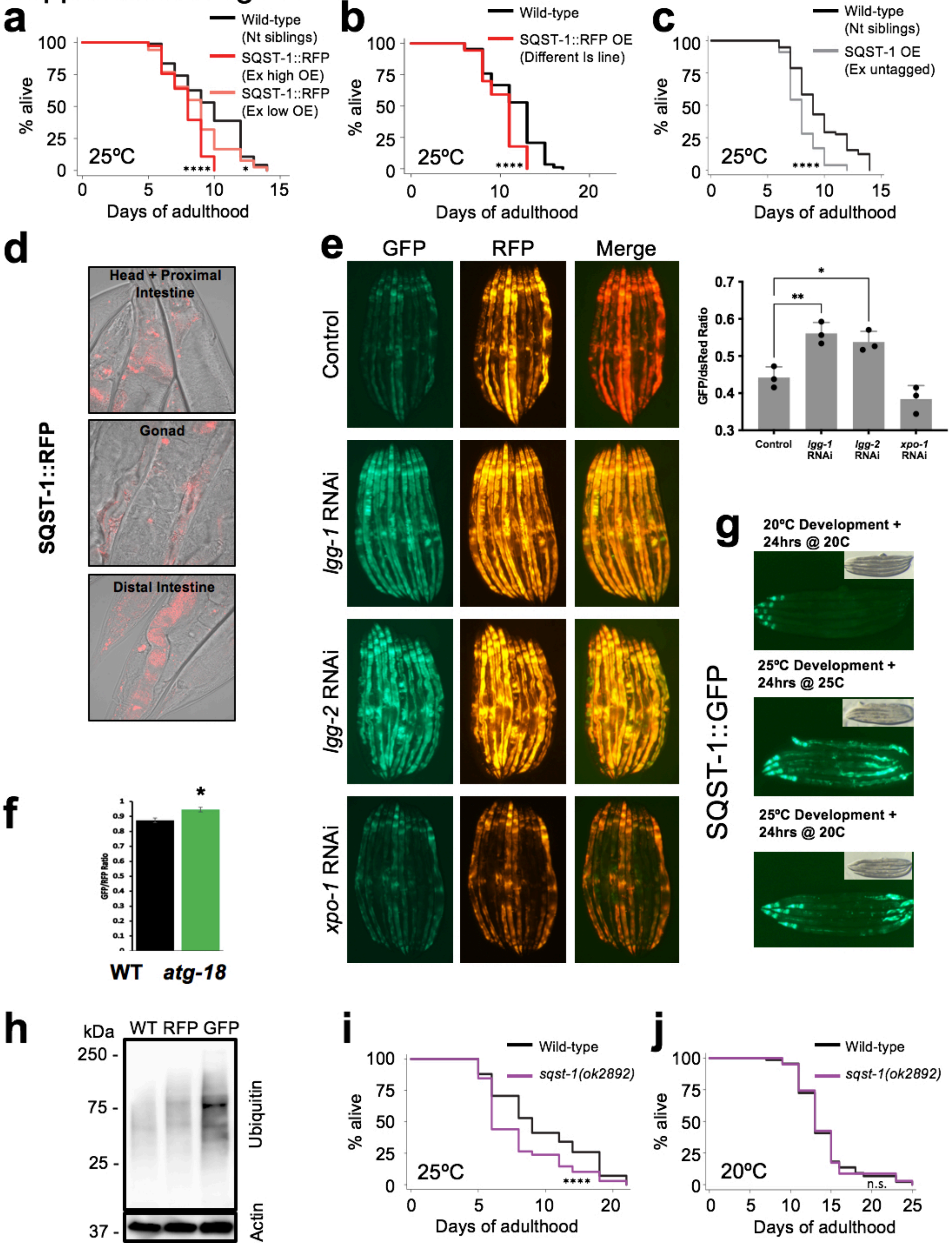

**Supplemental Figure 1. SQST-1 function in lifespan modulation is temperature-dependent. a.** Lifespan analysis of wild-type animals (non-transgenic siblings) and transgenic animals over-expressing SQST-1::RFP at low and high level (Ex: Extrachromosomal array), and **b.** transgenic animals over-expressing SQST-1::RFP (Integrated strain, Is) raised at 20ºC and grown at 25ºC during adulthood on OP50 *E. coli*. n=100. **c.** Lifespan analysis of wild-type animals (non-transgenic siblings, Nt) and transgenic animals over-expressing SQST-1 (untagged). **d.** Confocal images of SQST-1::RFP-expressing animals grown at 25ºC for 5 days. **e.** Micrographs of animals expressing SQST-1::GFP::RFP after 72 hours of feeding during adulthood control bacteria or bacteria expressing dsRNA against autophagy genes *lgg-1* or *lgg-2,* or nuclear export protein and HLH-30 modulator *xpo-1*. **f.** Quantification of the GFP and RFP signal ratio in GFP::RFP::SQST-1 over-expressing animals in a wild-type or autophagy-defective *atg-18(gk378)* background. Average of 10 worms per image, n=4 images per condition *t-test* **p*<0.05, ***p*<0.01. **g.** Images (GFP and transmitted light) of animals over-expressing SQST-1::GFP were raised at 20ºC or 25ºC and kept or transferred to 20ºC, or kept at 25ºC for 24 hours. **h.** Immunoblot of ubiquitin and actin in Day 1 wild-type (WT) animals and SQST-1::RFP (RFP) and SQST-1::GFP (GFP) over-expressing animals raised at 25ºC. Lifespan analysis of wild-type animals and *sqst-1(ok2892)* animals raised at 20ºC and grown at 25ºC (**i**) or 20ºC (**j**) during adulthood on OP50 *E. coli*. n=100. Details on lifespan analyses and repeats are available in Supplemental Table 3, Mantel-Cox log-rank. n.s.: not significant, **p<0.05,****p<0.001,*.

### Supplemental Figure 2

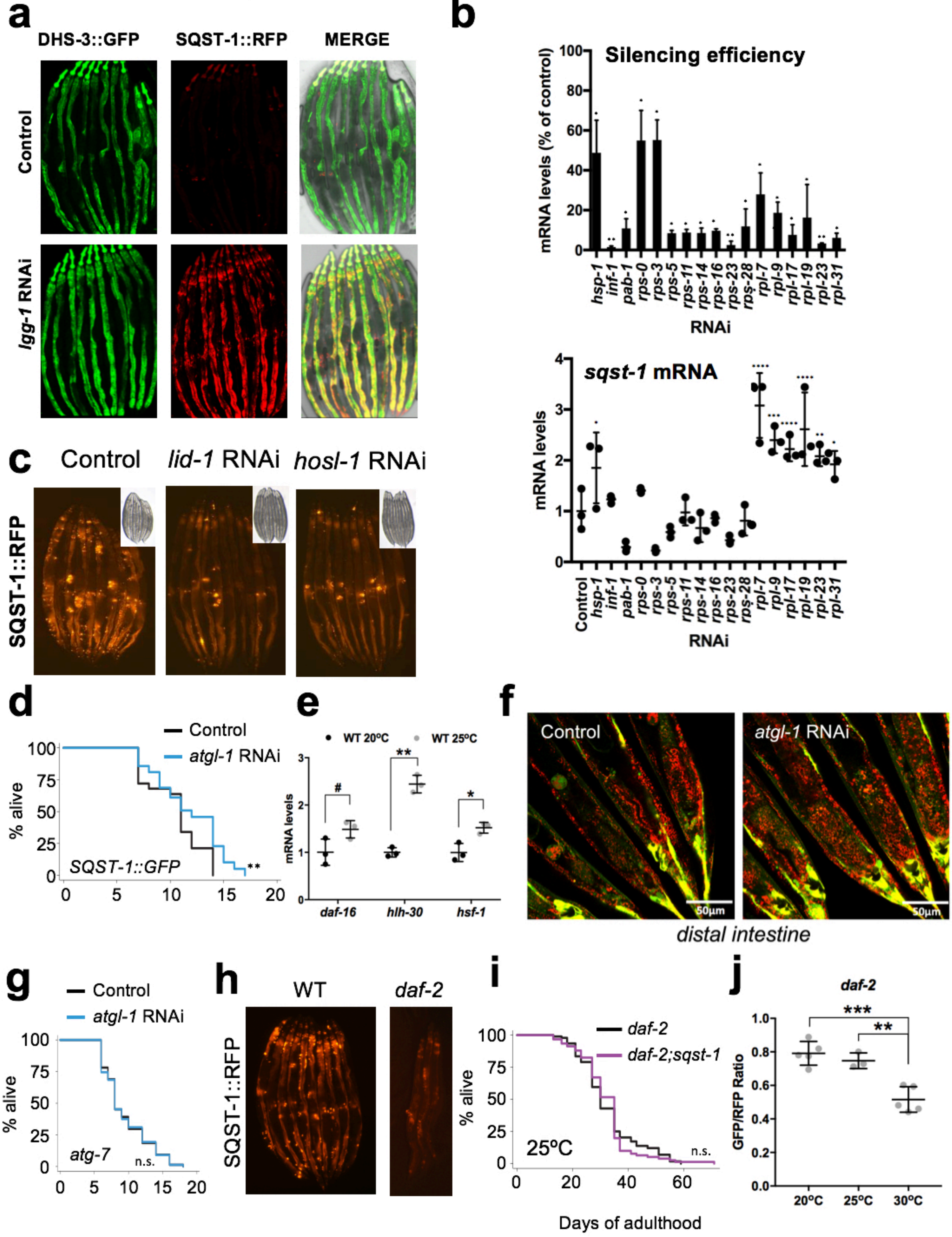

**Supplemental Figure 2. Lipid droplets enhance lifespan, autophagy, and the response to heat stress a.** Image of animals expressing both SQST-1::RFP and the lipid droplet-resident protein DHS-3 fused to GFP, subjected to *lgg-1* silencing for 3 days during adulthood. **b.** qPCR analysis of gene silencing efficiency of each modifier and their corresponding levels of *sqst-1* mRNA after 4 days of silencing during adulthood of animals expressing SQST-1::RFP. n=3, ±SD, *t-test* (silencing efficiency) or *ANOVA* (*sqst-1* mRNA) *$p<0.05$, **$p<0.01$. **d.** Lifespan analysis of transgenic animals over-expressing SQST-1::GFP grown on control bacteria or bacteria expressing dsRNA against *atgl-1*. **e.** qPCR analysis *daf-16, hlh-30* and *hsf-1* in Day 1 wild-type animals raised at 20ºC or 25ºC on OP50 *E. coli*. Biological triplicates *t-test* #$p=0.06$, *$p<0.05$, **$p<0.01$ **f.** Representative confocal microscopy images of the distal intestine of animals expressing mCherry::GFP::LGG-1 and raised at 20ºC and fed during adulthood control bacteria or bacteria expressing dsRNA against *atgl-1* for 2 days at 25ºC. **g.** Lifespan analysis of *atg-7(bp411)* mutants fed control bacteria or bacteria expressing dsRNA against *atgl-1*. **h.** Image of *daf-2(e1370)* animals over-expressing SQST-1::RFP raised on OP50 *E. coli*. bacteria at 20ºC and transferred to 25ºC during adulthood for 5 days. Comparative WT image in Figure 1a. **i.** Lifespan analysis of *daf-2(e1370)* and *daf-2(e1370);sqst-1(ok2892)* raised at 20ºC and grown at 25ºC during adulthood on OP50 *E. coli*. bacteria (n=100). **j.** Levels of GFP and RFP were measured in transgenic tandem *daf-2;SQST-1::GFP::RFP* animals after incubating Day 1 animals at 20°C, 25°C or 30°C for 24 hours on OP50 *E. coli*. Average of 10 worms per image. n=3-5 image per condition *t-test* **$p<0.01$, ***$p<0.001$. Details on lifespan analyses and repeats are available in Supplemental Tables 3 and 4, Mantel-Cox log-rank. n.s.: not significant, **$p<0.01$.

### Supplemental Figure 3

**a**

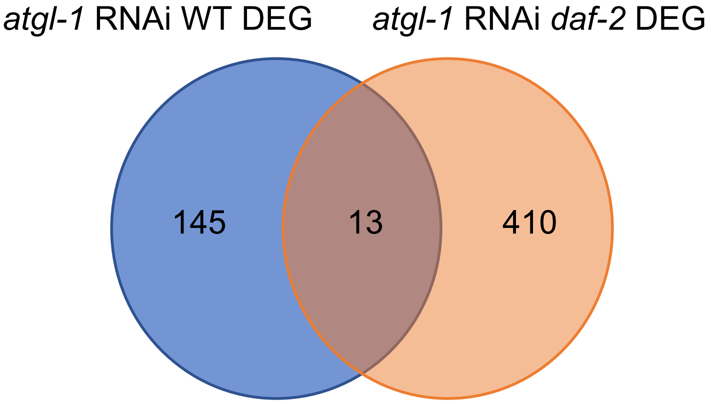


**b**

| *Gene Name* | Regulation in WT on *atgl-1* RNAi | Regulation in *daf-2* on *atgl-1* RNAi |
|---|---|---|
| ***atgl-1*** | Down | Down |
| *col-130* | Down | Up |
| *Y47D7A.13* | Down | Down |
| *nhx-2* | Down | Up |
| *T19B10.2* | Down | Down |
| *col-179* | Down | Down |
| *W03F9.4* | Down | Down |
| *ttr-23* | Up | Down |
| *mup-4* | Down | Down |
| *col-8* | Down | Up |
| *ttr-26* | Up | Down |
| *pvf-1* | Up | Up |
| *col-124* | Down | Down |

**c**

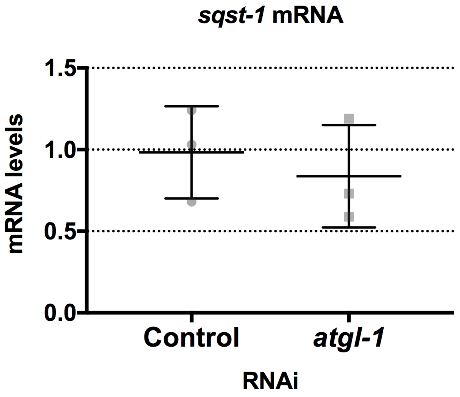


**d**

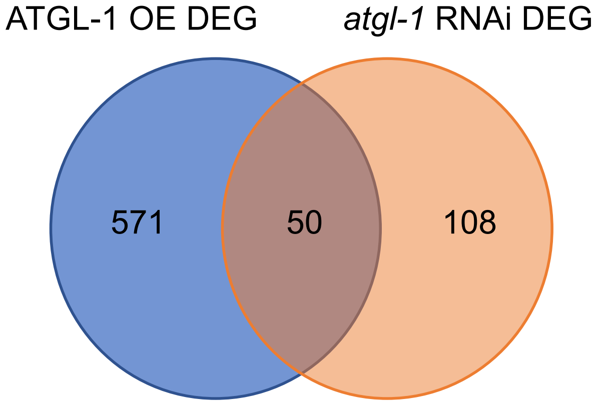


**e**

| *Gene Name* | Regulation in ATGL-1 OE | Regulation in *atgl-1* RNAi |
|---|---|---|
| ***atgl-1*** | Up | Down |
| *C55A6.7* | Down | Down |
| *F10D2.10* | Up | Down |
| *col-130* | Down | Down |
| *cest-1* | Down | Down |
| *ifp-1* | Down | Down |
| *T19B10.2* | Up | Down |
| *pcp-3* | Down | Down |
| *col-14* | Down | Down |
| *grl-16* | Down | Down |
| *mlc-1* | Down | Down |
| *lys-10* | Down | Down |
| *W03F9.4* | Down | Down |
| *aldo-1* | Down | Down |
| *col-104* | Down | Down |
| *col-77* | Down | Down |
| *gsto-1* | Down | Down |
| *wrt-6* | Down | Down |
| *C01B9.1* | Down | Down |
| *T05E7.1* | Down | Down |
| *clec-47* | Up | Down |
| *npa-1* | Down | Down |
| *M04C3.2* | Down | Down |
| *ttr-26* | Up | Up |
| *T19D12.1* | Down | Down |
| *col-125* | Down | Down |
| *dim-1* | Down | Down |
| *ifc-1* | Down | Down |
| *col-97* | Down | Down |
| *M153.1* | Down | Down |
| *poml-3* | Down | Down |
| *W08A12.2* | Up | Up |
| *pvf-1* | Up | Up |
| *M153.2* | Down | Down |
| *Y16B4A.2* | Down | Down |
| *acs-7* | Down | Down |
| *F10D2.8* | Down | Down |
| *H02F09.3* | Down | Down |
| *wrt-4* | Down | Down |
| *grd-6* | Down | Down |
| *idhg-2* | Down | Down |
| *cyp-14A2* | Down | Down |
| *ifd-1* | Down | Down |
| *Y43F8B.1* | Down | Down |
| *col-157* | Down | Down |
| *haf-9* | Down | Down |
| *ifc-2* | Down | Down |
| *T23E7.2* | Down | Down |
| *H43E16.1* | Down | Down |
| *tnt-2* | Down | Down |

**Supplemental Figure 3. Transcriptomic analyses of animals with over-expressed or silenced *atgl-1* reveal limited transcriptional changes.**

**a.** Overlap of differentially expressed genes (DEGs) in WT and *daf-2* mutant backgrounds when *atgl-1* is silenced during adulthood at 25ºC for 4 days. **b.** Identity of overlapping DEGs from a. and their regulation status in each strain. **c.** qPCR analysis of *sqst-1* mRNA in wild-type animals fed during adulthood control bacteria or bacteria expressing dsRNA against *atgl-1* for 4 days at 25ºC. n=3 ±SD *t*-test. **d.** Overlap of DEGs between nematodes over-expressing ATGL-1::GFP and adult-only *atgl-1* silencing in wild-type animals at 25ºC for 4 days. **e.** Identity of overlapping DEGs from c**.** and their regulation status under each condition.

### Supplemental Figure 4

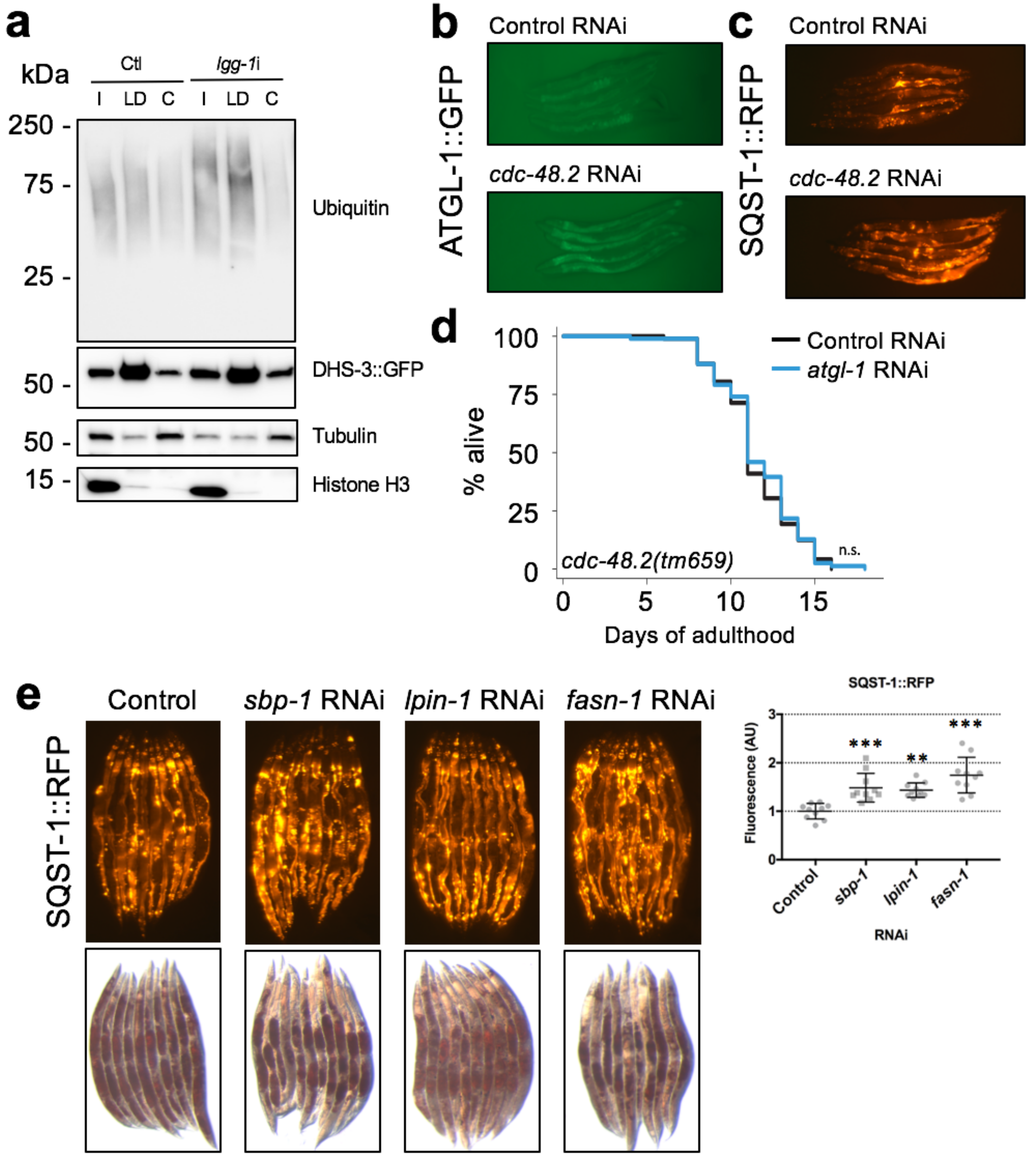

**Supplemental Figure 4. SQST-1 levels are modulated by CDC-48 activity and lipid droplet levels. a.** *daf-2* animals expressing lipid droplet-resident protein DHS-3 fused to GFP were raised at 20°C and then grown at 25°C during adulthood on control bacteria or bacteria expressing RNAi against *lgg-1* for 4 days. Levels of ubiquitinated proteins, DHS-3::GFP, cytosolic marker tubulin and nuclear marker histone H3 were immunoblotted from total input (I), cytosol (C) and lipid droplet (LD) fractions (fractions loaded comparatively, i.e. 10%). **b.** Day 1 ATGL-1::GFP and **c.** SQST-1::RFP over-expressing nematodes were fed control bacteria or bacteria expressing dsRNA against *cdc-48.2* for 2 days at 25ºC. Lifespan analysis of **d.** *cdc-48.2* mutants fed control bacteria or bacteria expressing dsRNA against *atgl-1* during adulthood at 25ºC. **e.** Day 1 SQST-1::RFP over-expressing nematodes were fed control bacteria or bacteria expressing dsRNA against *sbp-1, lpin-1,* or *fasn-1* for 6 days at 25ºC. Fluorescence was quantified using Image J. ±SD *ANOVA* *$p<0.05$,****$p<0.001$*. Corresponding images of nematodes stained with Oil-Red-O. Details on lifespan analyses and repeats are available in Supplemental Table 4, Mantel-Cox log-rank. n.s.: not significant.

### Supplemental Table 1

**See Supplemental Table 1 Excel File**

### Supplemental Table 2

**See Supplemental Table 2 Excel File**

### Supplemental Table 3

| Strains | Transgenic Mean Lifespan | Events Observed | N2 Control Mean Lifespan | Events Observed | % Difference | P Value | Fig. |
|---|---|---|---|---|---|---|---|
| **25ºC** | | | | | | | |
| *LRL160 (llcIs12 (sqst-1p::sqst-1::RFP::unc-54 + myo-2p::GFP))* | 11.9 | 43/100 | 13.5 | 50/100 | -11% | 0.0661 | |
| *LRL160 (llcIs12 (sqst-1p::sqst-1::RFP::unc-54 + myo-2p::GFP))* | 10.5 | 76/100 | 11.5 | 87/100 | -9% | 0.0094 | |
| *LRL160 (llcIs12 (sqst-1p::sqst-1::RFP::unc-54 + myo-2p::GFP))* | 9.8 | 57/100 | 12.9 | 49/100 | -24% | <0.0001 | 1a |
| *LRL161 (llcIs13 (sqst-1p::sqst-1::RFP::unc-54 + myo-2p::GFP))* | 10.1 | 84/100 | 11.5 | 87/100 | -12% | <0.0001 | S1b |
| *MAH349(sqIs35[pMH951/SQST::GFP/p62::GFP+pMH876/unc-122p::rfp])* | 7.7 | 50/100 | 9.9 | 53/100 | -22% | <0.0001 | |
| *MAH349(sqIs35[pMH951/SQST::GFP/p62::GFP+pMH876/unc-122p::rfp])* | 7.9 | 48/100 | 10.0 | 86/100 | -21% | <0.0001 | |
| *MAH349(sqIs35[pMH951/SQST::GFP/p62::GFP+pMH876/unc-122p::rfp])* | 7.9 | 58/100 | 10.6 | 71/100 | -26% | <0.0001 | 1b |
| *MAH349(sqIs35[pMH951/SQST::GFP/p62::GFP+pMH876/unc-122p::rfp])* | 9.6 | 28/100 | 11.0 | 48/100 | -13% | 0.0320 | |
| *LRL132 (llcEx55 (sqst-1p::sqst-1::GFP::RFP::unc-54))* | 10.7 | 89/100 | 11.7 | 74/100 | -9% | 0.0017 | 1c |
| *LRL132 (llcEx55 (sqst-1p::sqst-1::GFP::RFP::unc-54))* | 10.0 | 89/100 | 10.9 | 70/100 | -8% | 0.0030 | |
| *LRL90( llcEx39 (sqst-1p::sqst-1::RFP::unc-54 + myo-2p::GFP)) Ex high* | 7.9 | 124/160 | 9.6 | 57/100 | -18% | <0.0001 | S1a |
| *LRL90( llcEx39 (sqst-1p::sqst-1::RFP::unc-54 + myo-2p::GFP)) Ex low* | 8.6 | 79/100 | 9.6 | 57/100 | -11% | 0.0250 | S1a |
| *LRL90( llcEx39 (sqst-1p::sqst-1::RFP::unc-54 + myo-2p::GFP)) Ex high* | 9.8 | 87/100 | 11.2 | 75/100 | -13% | 0.0061 | |
| *LRL90( llcEx39 (sqst-1p::sqst-1::RFP::unc-54 + myo-2p::GFP)) Ex low* | 9.5 | 69/100 | 11.2 | 75/100 | -15% | 0.0021 | |
| *MAH844 (Exsq146 (sqst-1p::sqst-1 + rol-6))* | 8.0 | 61/100 | 9.6 | 68/100 | -16% | <0.0001 | S1c |
| *VC2196 (sqst-1 (ok2892) IV)* | 7.5 | 87/100 | 9.9 | 53/100 | -24% | <0.0001 | |
| *VC2196 (sqst-1 (ok2892) IV)* | 7.9 | 81/100 | 10.0 | 86/100 | -21% | <0.0001 | S1i |
| *VC2196 (sqst-1 (ok2892) IV)* | 9.3 | 75/100 | 10.6 | 71/100 | -12% | 0.0420 | |
| *VC2196 (sqst-1 (ok2892) IV)* | 9.5 | 63/100 | 11.0 | 48/100 | -14% | 0.0023 | |
| *VS20 (hjIs67 [atgl-1p::atgl-1::GFP + mec-7::RFP])* | 11.8 | 74/100 | 13.5 | 50/100 | -12% | 0.0203 | 4a |
| *VS20 (hjIs67 [atgl-1p::atgl-1::GFP + mec-7::RFP])* | 10.1 | 59/100 | 10.7 | 79/100 | -6% | 0.0312 | |
| *VS20 (hjIs67 [atgl-1p::atgl-1::GFP + mec-7::RFP])* | 11.6 | 61/100 | 12.9 | 49/100 | -10% | 0.0035 | |
| *LRL24 (daf-2(e1370) III; sqIs35[pMH951/SQST::GFP/p62::GFP+pMH876/unc-122p::rfp])* | 30.7 | 95/100 | 27.9 (*daf-2*) | 95/100 | 10% | 0.1258 | |
| *LRL24 (daf-2(e1370) III; sqIs35[pMH951/SQST::GFP/p62::GFP+pMH876/unc-122p::rfp])* | 31.9 | 92/100 | 29.3 (*daf-2*) | 89/100 | 9% | 0.5998 | 3j |
| *LRL170 (daf-2(e1370) III; sqst-1 (ok2892) IV)* | 32.0 | 89/100 | 32.5 (*daf-2*) | 80/100 | -2% | 0.8749 | S2e |
| **20ºC** | | | | | | | |
| *LRL160 (llcIs12 (sqst-1p::sqst-1::RFP::unc-54 + myo-2p::GFP))* | 14.0 | 62/100 | 15.1 | 68/100 | -7% | 0.1021 | 1d |
| *LRL160 (llcIs12 (sqst-1p::sqst-1::RFP::unc-54 + myo-2p::GFP))* | 15.3 | 48/100 | 16.3 | 50/100 | -6% | 0.2725 | |
| *LRL161 (llcIs13 (sqst-1p::sqst-1::RFP::unc-54 + myo-2p::GFP))* | 14.0 | 63/100 | 15.1 | 68/100 | -7% | 0.0904 | |
| *MAH349(sqIs35[pMH951/SQST::GFP/p62::GFP+pMH876/unc-122p::rfp])* | 14.1 | 42/100 | 14.0 | 49/100 | 1% | 0.6271 | 1e |
| *MAH349(sqIs35[pMH951/SQST::GFP/p62::GFP+pMH876/unc-122p::rfp])* | 15.3 | 48/100 | 14.2 | 30/100 | 7% | 0.3137 | |
| *LRL132 (llcEx55 (sqst-1p::sqst-1::GFP::RFP::unc-54))* | 14.1 | 68/100 | 15.0 | 42/100 | -6% | 0.1409 | 1f |
| *VC2196 (sqst-1 (ok2892) IV)* | 14.1 | 52/100 | 14.0 | 49/100 | 1% | 0.8730 | S1j |
| *VS20 (hjIs67 [atgl-1p::atgl-1::GFP + mec-7::RFP])* | 15.2 | 77/100 | 11.9 | 58/100 | 27% | <0.0001 | |

**Supplemental Table 3. Lifespan analyses performed on OP50 *E. coli*.** Animals were raised at 20ºC and grown at 20ºC or 25ºC (as noted) during adulthood on OP50 *E. coli*. Mantel-Cox log-rank.

### Supplemental Table 4

| Strains | *atgl-1* RNAi Mean Lifespan | Events Observed | Control RNAi Mean Lifespan | Events Observed | % Difference | P Value | Fig. |
|---|---|---|---|---|---|---|---|
| ***N2*** | 16.0 | 59/100 | 13.1 | 53/100 | 22% | 0.0017 | 3a |
| ***N2*** | 11.9 | 75/100 | 10.4 | 79/100 | 14% | 0.0019 | |
| ***N2*** | 12.6 | 90/100 | 10.6 | 79/100 | 19% | <0.0001 | |
| ***N2*** | 12.7 | 66/100 | 9.9 | 50/100 | 28% | <0.0001 | |
| ***N2*** | 11.4 | 52/100 | 10.2 | 60/100 | 12% | 0.0461 | |
| ***N2*** | 11.5 | 64/100 | 10.1 | 72/100 | 14% | 0.0009 | |
| ***N2*** | 11.8 | 57/100 | 9.9 | 67/100 | 19% | 0.0001 | |
| ***N2*** | 13.1 | 75/100 | 11.1 | 53/100 | 18% | 0.0231 | |
| ***N2*** | 12.7 | 62/100 | 10.5 | 61/100 | 21% | <0.0001 | |
| ***N2*** | 12.8 | 55/100 | 11.7 | 65/100 | 9% | 0.0468 | |
| ***CF1037 (daf-16 (mu86) I)*** | 14.0 | 55/100 | 11.6 | 53/100 | 21% (22) | 0.0010 | 3e |
| ***CF1037 (daf-16 (mu86) I)*** | 10.1 | 83/100 | 9.0 | 81/100 | 12% (14) | 0.0002 | |
| ***LRL31 (hlh-30 (tm1978) IV)*** | 14.7 | 72/100 | 12.1 | 62/100 | 22% (22) | <0.0001 | 3f |
| ***LRL31 (hlh-30 (tm1978) IV)*** | 8.7 | 59/100 | 7.9 | 77/100 | 10% (14) | <0.0001 | |
| ***CF2495 (hsf-1 (sy441) I)*** | 12.4 | 36/100 | 11.9 | 35/100 | 4% (22) | 0.5855 | 3g |
| ***CF2495 (hsf-1 (sy441) I)*** | 10.7 | 86/100 | 10.2 | 85/100 | 5% (14) | 0.3276 | |
| ***MAH349 (sqls35[sqst-1p::sqst-1::GFP + unc-122p::rfp])*** | 11.8 | 40/100 | 9.8 | 44/100 | 20%(19) | 0.0010 | S2d |
| ***LRL160 (llcls12 (sqst-1p::sqst-1::RFP::unc-54 + myo-2p::GFP))*** | 12.1 | 73/100 | 8.6 | 69/100 | 41% (19) | <0.0001 | 3c |
| ***HZ1686 (atg-7 bp411 IV)*** | 9.5 | 85/100 | 9.5 | 82/100 | 0%(14) | 0.9579 | S2g |
| ***HZ1686 (atg-7 bp411 IV)*** | 9.0 | 73/100 | 9.3 | 58/100 | -3% (19) | 0.5323 | |
| ***CF1041 (daf-2 e1370 III)*** | 35.0 | 80/100 | 27.3 | 76/100 | 28% (18) | <0.0001 | 3j |
| ***CF1041 (daf-2 e1370 III)*** | 25.1 | 60/100 | 23.0 | 81/100 | 9% (18) | 0.0312 | |
| ***FX544 (cdc-48.1 (tm544) II)*** | 12.4 | 79/100 | 12.3 | 77/100 | -1% (21) | 0.7901 | |
| ***FX659 (cdc-48.2 (tm659) II)*** | 9.4 | 47/100 | 9.5 | 54/100 | 1% (12) | 0.8290 | S4d |
| ***FX659 (cdc-48.2 (tm659) II)*** | 11.6 | 79/100 | 11.5 | 76/100 | 1% (21) | 0.5981 | |

**Supplemental Table 4. Lifespan analyses related to *atgl-1* silencing.** Animals were developed at 20ºC on OP50 and transferred on control bacteria or bacteria expressing dsRNA against *atgl-1* and grown at 25ºC during adulthood. Mantel-Cox log-rank.

### Supplemental Table 5

| Published strains used in this study: | | |
|---|---|---|
| **Strain** | **Genotype** | **Strain origin** |
| **N2** | Wild-type, WT | Hansen Lab |
| **CF1037** | *daf-16 (mu86) I* | Hansen Lab |
| **CF1041** | *daf-2(e1370) III* | Hansen Lab |
| **CF2495** | *hsf-1(sy441) I* | Hansen Lab |
| **FX544** | *cdc-48.1(tm544) II* | CGC, National Bioresource Project at the Tokyo Women's Medical University School of Medicine |
| **FX659** | *cdc-48.2(tm659) II* | CGC, National Bioresource Project at the Tokyo Women's Medical University School of Medicine |
| **FX1978** | *hlh-30 (tm1978) IV* | CGC, National Bioresource Project at the Tokyo Women's Medical University School of Medicine |
| **GF80** | dgEx80 [(pAMS66) vha-6p::Q44::YFP + rol-6(su1006) + pBluescript II]. | CGC |
| **GMC101** | dvIs100 [unc-54p::A-beta-1-42::unc-54 3'-UTR + mtl-2p::GFP] | CGC |
| **LIU1** | *ldrIs1 (dhs-3p::dhs-3::GFP + unc-76(+))* | CGC |
| **LRL9** | *atg-7 (bp411) IV* | Lapierre Lab |
| **MAH78** | *sqIs2[Plipl-4::LIPL-4(K04A8.5)::SL2gfp + Pmyo-2::CHERRY]* | Hansen Lab |
| **MAH215** | *sqIs15[lgg-1p::mCherry::GFP::lgg-1 + rol-6]* | Hansen Lab |
| **MAH349** | *sqIs35[sqst-1p::sqst-1::GFP + unc-122p::rfp]* | Hansen Lab |
| **MAH844** | *sqEx146[sqst-1p::sqst-1 + rol-6]* | CGC |
| **VC2196** | *sqst-1 (ok2892) IV* | CGC |
| **VS20** | *hjIs67 [atgl-1p::atgl-1::GFP + mec-7::RFP]* | CGC |

**Supplemental Table 5. Published strains used in the study**

### Supplemental Table 6

| New strains created for this study: | | |
|---|---|---|
| **Strain** | **Genotype** | **Comments** |
| **LRL24** | *daf-2(e1370) III; sqIs35[pMH951/SQST ::GFP/p62::GFP+pMH876/unc-122p::rfp]* | CF1041 x MAH349 |
| **LRL31** | *hlh-30 (tm1978) IV* | FX1978 4x backcrossed to N2 |
| **LRL90** | *llcEx39 (sqst-1p::sqst-1::RFP::unc-54 + myo-2p::GFP)* | Injected N2 with 20 ng/uL pLAP26 and 5 ng/uL pLP7. pLP7 (*myo-2p::GFP::unc-54*) was obtained from Addgene (pBCN27). pLAP26 *(sqst-1p::sqst-1::RFP::unc-54)* was generated by NEBuilder® HiFi DNA Assembly Cloning Kit (New England BioLabs Inc., Ipswich, MA) by replacing the *3XFLAG*-tag with the *RFP* sequence in pLP25 *(sqst-1p::sqst-1::3XFLAG::unc-54)* (unpublished). The *RFP* insert was PCR-amplified from Addgene plasmid #8938 (*unc-122p::RFP)* with HiFi-compatible forward primer LC209 (5' GATGTGTCTTCAGGCGCTTCTTCACGGCTCGGGCTCGATGGTGCGCTCCTCCAAGAACG 3') and reverse primer LC210 (5' GACACCAGACAAGTTGGTAATGGCTACAGGAACAGGTGGTGGC 3'). For linearizing and amplifying linear fragments of the vector backbone *sqst-1p::sqst-1::unc-54* from pLAP25, HiFi-compatible forward primer LC207 (5' CCATTACCAACTTGTCTGGTGTC 3') and reverse primer LC208 (5' GTGAAGAAGCGCCTGAAGACACATC 3') were used. Ligation of insert to the vector backbone and subsequent transformation of products into competent *E. coli*, were performed according to manufacturer's instructions. |
| **LRL132** | *llcEx55 (sqst-1p::sqst-1::GFP::RFP::unc-54)* | Injected N2 with 20 ng/uL pLAP29 *(sqst-1p::sqst-1::GFP::RFP::unc-54).* pLAP29 was generated by NEBuilder® HiFi DNA Assembly Cloning Kit (New England BioLabs Inc., Ipswich, MA) by inserting the *GFP* sequence 3' and 5' of the *sqst-1* and *RFP* sequences in pLAP26 *(sqst-1p::sqst-1::RFP::unc-54)* respectively. The *GFP* insert was PCR-amplified from plasmid #836 kindly provided by Dr Andrew Dillin (UC Berkeley) with HiFi-compatible forward primer LC251 (5' TTCAGGCGCTTCTTCACGGCTCGGGCTCGATGAGTAAAGGAGAAGAACTTTTC 3') and reverse primer LC252 (5' GACGTTCTTGGAGGAGCGCACCATCGAGCCGCCGCCGCCTTTGTATAGTTCATCCATGCCATG 3'). For linearizing and amplifying the vector backbone pLP26, HiFi-compatible forward primer LC244 (5' GATGGTGCGCTCCTCCAAGAACGTC 3') and reverse primer LC250 (5' GCCGTGAAGAAGCGCCTGAAGACAC 3') were used. Ligation of insert to the vector backbone and subsequent transformation of products into competent *E. coli*, were performed according to manufacturer's instructions. |
| **LRL160** | *llcIs12 (sqst-1p::sqst-1::RFP::unc-54 + myo-2p::GFP)* | UV irradiation of LRL90, 6x backcrossed to N2 |
| **LRL161** | *llcIs13 (sqst-1p::sqst-1::RFP::unc-54 + myo-2p::GFP)* | UV irradiation of LRL90, 6x backcrossed to N2 |
| **LRL170** | *daf-2(e1370) III; sqst-1 (ok2892) IV* | CF1041 x VC2196 |
| **LRL171** | *daf-2(e1370) III; llcIs12 (p62p::p62::RFP ::unc-54 + myo-2p::GFP)* | CF1041 x LRL160 |
| **LRL175** | *ldrIs1 (dhs-3p::dhs-3::GFP + unc-76(+)); llcIs12 (sqst-1p::sqst-1::RFP::unc-54 + myo-2p::GFP)* | LRL87 (LIU1 4x backcrossed into N2) x LRL160 |
| **LRL178** | *hjIs67 [atgl-1p::atgl-1::GFP + mec-7::RFP]; llcIs12 (sqst-1p::sqst-1::RFP::unc-54 + myo-2p::GFP)* | VS20 x LRL160 |

#### Supplemental Table 6. New strains created for this study

Supplemental Table 7

| Genes | Direction | Sequence |
|---|---|---|
| *act-1* | Forward | CTACGAACTTCCTGACGGACAAG |
| *act-1* | Reverse | CCGGCGGACTCCATACC |
| *atgl-1* | Forward | CCGGACGTCTGGTTATCTCG |
| *atgl-1* | Reverse | GCTCGTCGTAGATTGGCTGA |
| *cyn-1* | Forward | GTGTCACCATGGAGTTGTTC |
| *cyn-1* | Reverse | TCCGTAGATTGATTCACCAC |
| *daf-16* | Forward | ATCCAATTGTGCCAAGCACTAA |
| *daf-16* | Reverse | CCACCATTTTGATAGTTTCCATAGG |
| *hlh-30* | Forward | CTCATCGGCCGGCGCTCATC |
| *hlh-30* | Reverse | AGAACGCGATGCGTGGTGGG |
| *hsf-1* | Forward | GCGGCTCCGTATAAGAATGCGACTAGGC |
| *hsf-1* | Reverse | TTAAACCAAATTAGGATCCGATGGACTTGGAGTAC |
| *hsp-1* | Forward | CGCTCAGACCTTCACAACCT |
| *hsp-1* | Reverse | TGGAAAGACGTCCCTTGTCG |
| *inf-1* | Forward | CGGCTTGATCGAGGGAAACTA |
| *inf-1* | Reverse | GCCATAACGAGAGCTTGCAC |
| *rpl-7* | Forward | TCAAGCGCAGAAAGCAGAGA |
| *rpl-7* | Reverse | ACGAAGACGGAGGATCTGGA |
| *rpl-9* | Forward | ACCTTCACCGTCAAGAACCG |
| *rpl-9* | Reverse | GAAGTGGGACACGACGAACG |
| *rpl-17* | Forward | CGGAAAACAGCACCAAGTCG |
| *rpl-17* | Reverse | GATCGAGGAGGAAGTCAGCG |
| *rpl-19* | Forward | GTTTGGCTTCGGCCGTATTG |
| *rpl-19* | Reverse | AGAGCTCGTGGTAAAGGTGC |
| *rpl-23* | Forward | CAACACCGGAGCCAAGAACTTG |
| *rpl-23* | Reverse | CGTTAGCAGCGATTCTTGGC |
| *rps-0* | Forward | GATGTCGTCGTTGTTTCGGC |
| *rps-0* | Reverse | CGGGAGATCTTTCCACGGAG |
| *rps-3* | Forward | GGCTGCCAATCAAAACGTGA |
| *rps-3* | Reverse | AACCTTCTCGGCGTAGAGCT |
| *rps-5* | Forward | GGCCGATAACTGGGGATCTG |
| *rps-5* | Reverse | GCTTCTTTCCGTTGTTGCGT |
| *rps-11* | Forward | CATTCGTGAGGTCGGACTCG |
| *rps-11* | Reverse | AAGCCCTTCTTGGAGGTTCC |
| *rps-14* | Forward | GGAATGAAGGTCAAGGCCGA |
| *rps-14* | Reverse | GCGACCTCCCTTTCTTCTGG |
| *rps-16* | Forward | GGTCGCCCACTTGAGTTCTT |
| *rps-16* | Reverse | TCCTGGTCCACCGAACTTCT |
| *rps-23* | Forward | GGAAAGCCGAAGGGACTCTG |
| *rps-23* | Reverse | GTCCGAAACCAGATACGAGCAC |
| *rps-28* | Forward | AGCTTACTCTTGCTCGTGTCA |
| *rps-28* | Reverse | AGTCTTCTGGCTTCTCTCTCAG |
| *sqst-1* | Forward | TGGCTGCTGCATCATCCGCT |
| *sqst-1* | Reverse | TCAATCGTGCCGAGACCGGG |

**Supplemental Table 7. qPCR primers sequences**
